# Systematic analysis of genetic vulnerabilities caused by chromosome loss events in cancer

**DOI:** 10.64898/2026.10.26.762060

**Authors:** Yi Bei, Jessica Keane, Mason Lynch, Kurt M. Tamaru, Suyeon Lee, Lu Qiao, Pranav K. Kandikuppa, Gregory Gauthier-Coles, Wenqi Zhao, Klaske M. Schukken, Ines Forrest, Sarah L. Thompson, Stephanie Roessler, Alison M. Taylor, Jason M. Sheltzer

**Affiliations:** Stanford University School of Medicine, Stanford, California 94305, USA; Yale University School of Medicine, New Haven, Connecticut 06511, USA; Heidelberg University, Medical Faculty, Institute of Pathology, University Hospital Heidelberg, 69120 Heidelberg, Germany; Columbia University School of Medicine, New York, New York 10032, USA

## Abstract

Chromosome loss events are common across tumor types but rarely occur in normal tissue, raising the possibility that these alterations could be exploited as cancer-specific therapeutic vulnerabilities. Here, we integrate genome-scale CRISPR screening data with chromosome-level copy number profiles from 1,150 cancer cell lines and identify 389 dependencies that are selectively associated with individual chromosome loss events. The most significant vulnerabilities uncovered by our analysis are paralogous gene pairs, in which the loss of a paralog on an aneuploid chromosome establishes a unique dependency on its redundant paralog encoded elsewhere in the genome. We experimentally validate two paralog dependencies: loss of chromosome 8p, which encodes the phosphatase *PPP2CB*, creates a specific dependency on its paralog *PPP2CA*, while loss of the Y chromosome, which encodes the helicase *DDX3Y*, creates a specific dependency on its paralog *DDX3X*. Systematic benchmarking of CRISPR screening data indicates that the selective dependency on DDX3X in loss-of-Y cells exceeds other published aneuploidy-associated vulnerabilities and matches or exceeds most clinically-validated biomarker-target relationships. Furthermore, we found that the acute degradation of DDX3X was sufficient to trigger apoptosis *in vitro* and tumor regressions in mice. Collectively, these findings establish chromosome loss as a recurrent source of biomarker-defined therapeutic targets, identify paralog insufficiency as a general vulnerability in aneuploid cells, and provide *in vivo* proof-of-concept for targeting DDX3X in cancers with Y chromosome loss.

## Introduction

Chromosome-scale copy number changes, otherwise known as aneuploidies, are found in approximately 90% of human cancers^1^. On a cellular level, aneuploidy alters the dosage of hundreds of genes at once, leading to the increased expression of oncogenes, reduced expression of tumor suppressor genes, and widespread disruption of cellular homeostasis^2–4^. In addition, chromosome loss events can result in loss of heterozygosity (LOH) for inactivating mutations in cancer genes, further amplifying the oncogenic consequences of aneuploidy^5,6^. Accordingly, the distribution of cancer aneuploidies is non-random: certain chromosomes are recurrently gained or lost across many different tumor types, independent of tissue of origin^7,8^. In The Cancer Genome Atlas (TCGA), the most commonly gained chromosomes include 8q, 7p, and 20q, which occur in 27% - 29% of all cancers^9^. In contrast, the most commonly lost chromosomes include 17p, 8p, and 18q, and these aneuploidies are found in 26% - 33% of all cancers^9^. Recently, loss of the Y chromosome (LOY) has been documented in ∼30% of cancers that arise in males, and this specific alteration has been linked with immune evasion, epigenetic plasticity, and aggressive disease^10–14^.

While aneuploidies are pervasive in cancer, single-cell sequencing studies have revealed that they are rarely observed in non-malignant tissue^15–18^. Thus, chromosome copy number alterations represent a prominent and pervasive genetic difference between cancer cells and normal cells. These observations raise the possibility that a therapy that selectively targets the aneuploid state could exhibit broad efficacy against multiple cancer types while also causing few side effects in healthy tissue.

Several genetic and pharmacological screens have been conducted in an attempt to identify therapeutic vulnerabilities that are conferred by aneuploidy^19–26^. These screens have identified certain genetic pathways that are preferentially-essential in cells harboring aneuploid chromosomes, including protein homeostasis^19,23^, DNA replication^24^, and the mitotic checkpoint machinery^21,27^. Relatedly, recent analyses have demonstrated that polyploid cells (e.g., cells that have undergone a whole-genome duplication and acquired an extra set of all chromosomes) require the kinesin KIF18A to successfully complete mitosis^21,27^. Based on these findings, multiple pharmaceutical companies have developed KIF18A inhibitors that are now undergoing patient testing^28^. These results provide proof-of-principle that changes in a cell’s ploidy can create druggable vulnerabilities, leading to the identification of new targets for clinical interventions.

To date, these attempts at identifying ploidy-associated targets have focused on chromosome gain events and gross changes in a cell’s karyotype. In contrast, vulnerabilities caused by specific chromosome loss events have not been comprehensively investigated. Nonetheless, chromosome losses exhibit several unique features that make them promising targets for therapeutic development: (1) Chromosome losses result in a significantly greater dosage change than chromosome gains (i.e., a two-fold change in expression vs. a 1.5-fold change in expression for an autosome, or, in the context of LOY, the complete loss of genes expressed from that chromosome). (2) While some degree of aneuploidy or chromosomal instability may be a consequence rather than a cause of tumorigenesis, increasing evidence has demonstrated that specific chromosome losses like 8p-loss and LOY are drivers of cancer development^6,11,29^. (3) Individual chromosome loss events represent a clear and unambiguous genomic lesion that can easily be identified by sequencing, FISH, or karyotyping as a patient-selection biomarker. For these reasons, we speculated that identifying dependencies that arise specifically from chromosome loss events could reveal a new set of selective, biomarker-driven therapeutic targets in cancer.

## Results

### Identification of genetic vulnerabilities associated with the loss of specific chromosomes

To uncover genetic vulnerabilities caused by specific chromosome loss events, we analyzed whole-genome CRISPR screening data from 1,150 cancer cell lines for which copy number data was available (Fig. 1A-B). We limited our analysis to 28 autosomal chromosome arms and the Y chromosome, each of which was lost in at least 150 cancer cell lines. First, we binned cell lines based on whether they had lost a copy of an individual chromosome or whether they were copy-neutral for that chromosome. We observed that groups of cell lines binned in this manner exhibited many additional chromosome copy number differences beyond the initial separation criteria. For instance, among cell lines that exhibited a loss of chromosome 8p, there were a median of 21 aneuploid chromosome arms, while cell lines that were neutral for chromosome 8p exhibited a median of 8 aneuploid chromosome arms (Fig. S1A). Similarly, cell lines that exhibited a loss of chromosome 8p were more likely to also harbor aneuploidies of 3p, 4p, 4q, and several other chromosomes compared to cell lines that were 8p-neutral. To address this discrepancy, we introduced an additional filtration step in our analysis, in which we eliminated cell lines with fewer than seven or more than 27 aneuploid chromosome arms. Applying these thresholds decreased the differences in both total and specific aneuploidy burden between cell lines separated based on individual aneuploidies (Fig. S1B-C).

**Figure 1.**
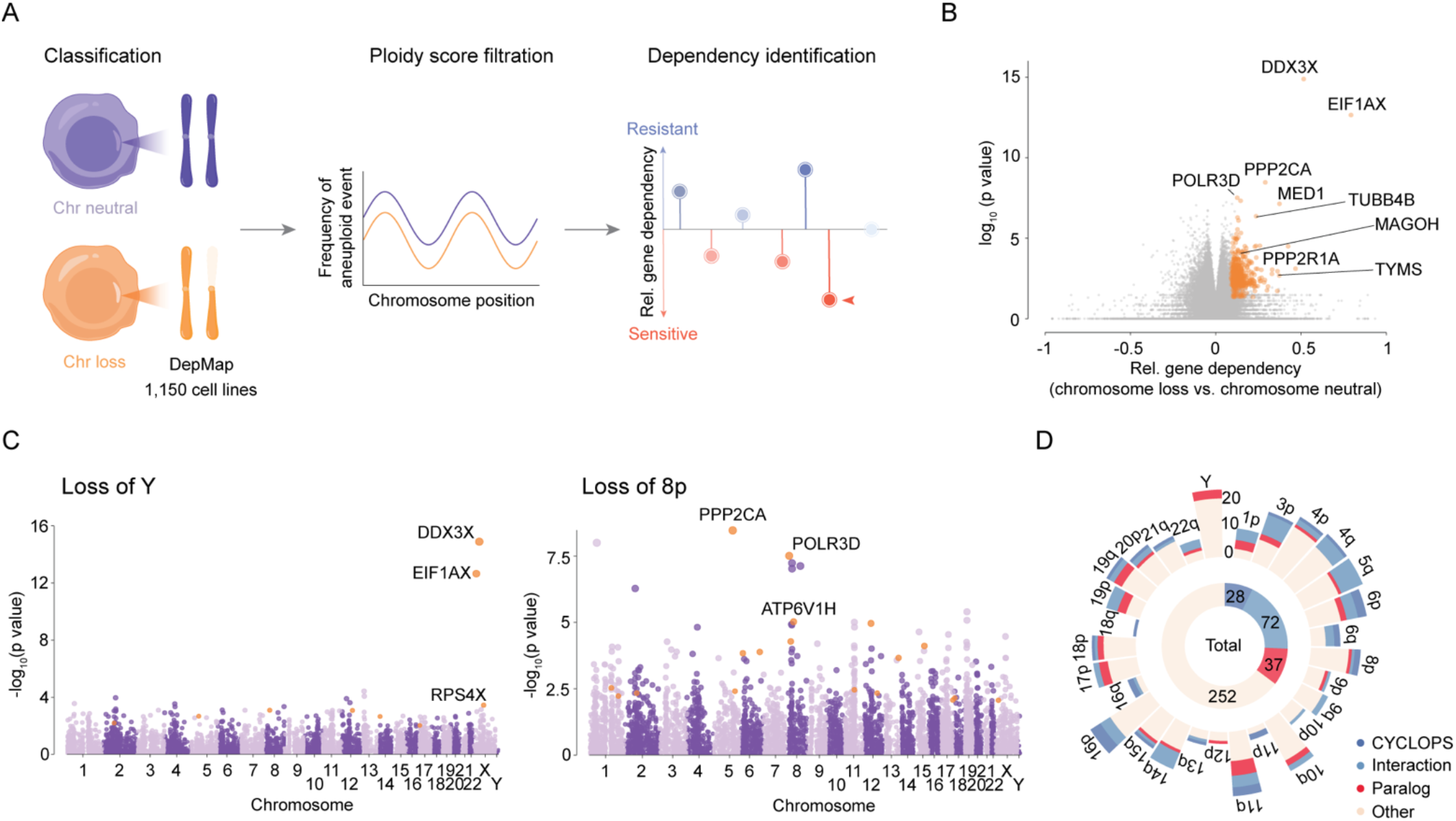
Chromosome loss events are associated with specific genetic dependencies. (A) Workflow for identifying chromosome-loss-associated dependencies in 1,150 cancer cell lines from DepMap. Cell lines were classified by chromosome copy-number status and filtered by aneuploidy score before comparing gene dependencies between chromosome-loss and chromosome-neutral cell lines. (B) Differential gene dependency across chromosome-loss events. Each point represents a gene–chromosome-loss association, plotted by relative dependency and statistical significance. Significant dependencies are highlighted in orange and selected genes are labeled. (C) Genome-wide analysis of differential gene dependency associated with Y chromosome loss (left) or chromosome 8p loss (right). Each point represents a gene, plotted according to its chromosomal location and the significance of the dependency difference between chromosome-loss and chromosome-neutral cell lines. Significant dependencies are highlighted in orange and selected genes are labeled. (D) Distribution of the identified dependencies across dependency classes. The outer stacked bars show the number of dependencies associated with each chromosome-arm loss, colored by class. The central doughnut chart summarizes all 389 dependencies: 28 CYCLOPS dependencies, 72 interactor dependencies, 37 paralog dependencies and 252 other dependencies.

Following this filtration step, we comprehensively identified all genes that exhibited a significant increase in essentiality in cells that had lost a specific chromosome. Two representative examples are shown in Figure 1C: the most significant vulnerabilities associated with loss of the Y chromosome were *DDX3X* and *EIF1AX*, while the most significant vulnerabilities associated with the loss of chromosome 8p were *PPP2CA* and *POLR3D*. Across all of the chromosomes included in this analysis, we identified four classes of significant vulnerabilities that were associated with specific chromosome loss events:

1. CYCLOPS genes (Copy-number alterations Yielding Cancer Liabilities Owing to Partial losS)^30^, in which the loss of a chromosome arm was associated with an increased sensitivity to the ablation of a gene encoded on that chromosome arm.
2. Paralog vulnerabilities, in which the loss of a chromosome arm was associated with an increased sensitivity to the ablation of a gene that is paralogous to a gene encoded on the lost chromosome arm.
3. Interaction vulnerabilities, in which the loss of a chromosome arm was associated with an increased sensitivity to the ablation of a gene whose protein product interacts with a gene encoded on the lost chromosome arm.
4. Other vulnerabilities, which could not be readily classified into one of the above categories (Fig. 1D).

In total, our comprehensive analysis uncovered 389 instances in which a gene exhibited a significant increase in essentiality in cells that had lost an individual chromosome arm (Table S1).

### Chromosome loss events are frequently associated with an increased sensitivity to paralog ablation

We noted that the three most-significant vulnerabilities and nine of the 20 most significant vulnerabilities across all chromosomes were paralogs of genes encoded on the lost chromosome (Fig. 2A-C and Table S1). We therefore decided to first focus on this class of aneuploidy-associated vulnerabilities. Two representative examples are shown in Fig. 2D-K: Loss of chromosome 8p, which encodes *PPP2CB*, was associated with an increased sensitivity to the ablation of its paralog *PPP2CA*, while loss of chromosome 11q, which encodes *PPP2R1B*, was associated with increased sensitivity to the ablation of its paralog *PPP2R1A*. In both cases, loss of the chromosome was associated with decreased expression of the paralog *in cis*, and we observed a highly-significant correlation across all cancer cell lines between the expression of the gene encoded on the aneuploid arm and sensitivity to the ablation of its paralog (Fig. 2F-G and Fig. 2J-K). We normalized and then combined all significant paralog vulnerabilities identified by our analysis, and we observed a consistent pattern in which chromosome losses decreased the expression of the paralog in *cis*, and decreasing expression of that paralog was significantly associated with increasing essentiality of the second paralog in *trans* (Fig. 2L-M). Considered together, these findings suggest that the reduced dosage of a gene resulting from chromosome loss increases the cellular dependence on its otherwise-redundant paralog.

**Figure 2.**
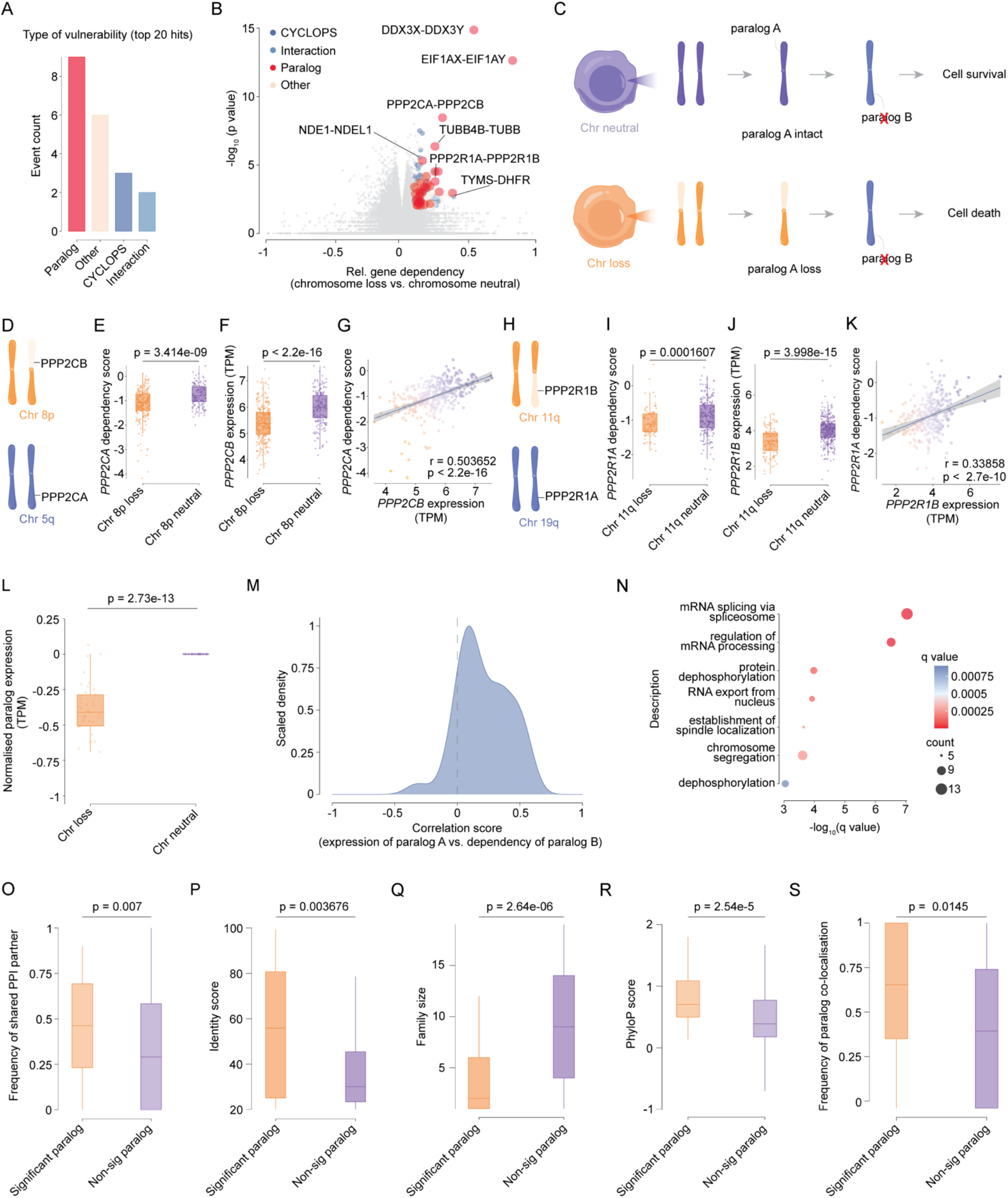
Paralog dependencies are the most frequent class among the top-ranked chromosome-loss-associated vulnerabilities. (A) Classification of the 20 top-ranked dependencies associated with specific chromosome loss events. (B) Differential gene dependency across chromosome-loss events, colored by dependency class: CYCLOPS, dark blue; interactor, light blue; paralog, red; and other, beige. The x-axis indicates relative dependency in chromosome-loss versus chromosome-neutral cells, and the y-axis indicates statistical significance (−log₁₀ P). Selected dependency–paralog pairs are labeled. (C) Model of a paralog dependency arising from chromosome loss. Disruption of one paralog is tolerated when its partner is retained, whereas reduced dosage of the partner following chromosome loss creates a dependency on the remaining paralog. (D) Chromosomal locations of *PPP2CB* on 8p and *PPP2CA* on 5q, illustrating the reduction in *PPP2CB* copy number following 8p loss. (E) *PPP2CA* dependency scores in cancer cell lines with chromosome 8p loss or neutral 8p copy-number status. (F) *PPP2CB* expression in cancer cell lines with chromosome 8p loss or neutral 8p copy-number status. (G) Relationship between *PPP2CB* expression and *PPP2CA* dependency scores. A fitted regression line is plotted against the data. (H) Chromosomal locations of *PPP2R1B* on 11q and *PPP2R1A* on 19q, illustrating the reduction in *PPP2R1B* copy number following 11q loss. (I) *PPP2R1A* dependency scores in cancer cell lines with chromosome 11q loss or neutral 11q copy-number status. (J) *PPP2R1B* expression in cancer cell lines with chromosome 11q loss or neutral 11q copy-number status. (K) Relationship between *PPP2R1B* expression and *PPP2R1A* dependency scores. A fitted regression line is plotted against the data. (L) Relative expression of paralogs located on lost chromosome arms, normalized to expression in the corresponding chromosome-neutral groups. (M) Distribution of correlation coefficients between expression of the paralog on the lost chromosome arm and dependency on its partner, across the 37 significant paralog pairs. The dashed line indicates a correlation coefficient of zero. (N) Gene ontology enrichment of significant paralog dependencies. Complete results are included in Table S2A. (O) Fraction of shared protein-protein interactions between significant and non-significant paralog pairs. (P) Sequence identity scores for significant and non-significant paralog pairs. (Q) Gene family sizes for significant and non-significant paralog pairs. (R) Evolutionary conservation (PhyloP score) for significant and non-significant paralog pairs. (S) Box plots comparing subcellular colocalization frequencies between significant and non-significant paralog pairs.

### Aneuploidy-associated paralog vulnerabilities cannot be explained by off-target CRISPR mutagenesis

As an alternate explanation for our results, we considered the possibility that the CRISPR gRNAs used to ablate one gene could exhibit promiscuous activity against their target’s paralogs. In a cell line monosomic for the chromosome encoding the paralog, only a single mutation of that gene would be required to eliminate its function, thereby sensitizing cells to this off-target activity and producing the apparent association between chromosome losses and paralog vulnerabilities. To address this possibility, we performed two analyses. First, we cloned 15 gRNAs targeting four significant aneuploidy-associated paralog vulnerabilities that were identified in our analysis (*FBXW11*, *PPP2CA*, *PPP2R1A*, and *TYMS*), using the same gRNA sequences and CRISPR expression system that were employed in the DepMap screens (Fig. S2A)^31^. We then transduced each gRNA into three different cell lines and performed deep sequencing to quantify CRISPR-induced mutations. While we could readily detect mutations in each gRNA’s target gene, we did not observe any increase in mutations in each gene’s paralog, indicating that our findings cannot be explained by off-target CRISPR activity across paralogs (Fig. S2B-C).

Second, we took advantage of the fact that most cancer cell lines included in our analysis had also been subjected to whole-genome shRNA screens, which typically target different sequences and exhibit different patterns of off-target activity compared to CRISPR screens^32^. We posited that if our findings were consistent across genetic screens conducted using both CRISPR and RNAi, then this would decrease the likelihood that they resulted from a CRISPR artifact. We re-analyzed the 10 most significant aneuploidy-associated paralog vulnerabilities identified in our initial CRISPR analysis using dependency data obtained from whole-genome shRNA screens, and we found that all 10 relationships remained significant in the RNAi analysis (Fig. S3). For instance, the RNA helicase *DDX17* is encoded on chromosome 22q, and we found that the loss of this chromosome was associated with an increased sensitivity to the ablation of its paralog *DDX5* in screens conducted using either RNAi or CRISPR (Fig. S3AH-AK). Moreover, the expression of the gene encoded on the lost chromosome was significantly correlated with increased sensitivity to RNAi-mediated ablation of its paralog across these top hits (Fig. S3). In total, these independent lines of evidence indicate that chromosome loss events sensitize cells to the ablation of specific paralogous genes, and these genetic relationships are readily detectable with both CRISPR and RNAi.

### Aneuploidy-associated paralog vulnerabilities are enriched in specific biological processes and are members of smaller gene families

We noted that our analysis identified 37 significant aneuploidy-associated paralog vulnerabilities, which represent only a small fraction of all paralogous genes encoded in the human genome. We therefore sought to understand the biological differences between these significant hits and the paralogous genes that were not identified as vulnerabilities in aneuploid cells. Gene ontology analysis of the aneuploidy-associated paralog vulnerabilities revealed a highly-significant enrichment of genes encoding splicing regulators (*DDX39A*, *HNRNPF*), protein phosphatases (*PPP2CA*, *PPP2R1A*), and chromosome segregation factors (*NDE1*, *DYNC1H1*)(Fig. 2N and Table S2A). Pairs of aneuploidy-associated paralog vulnerabilities exhibited higher identity scores and shared more protein-protein interactions with one-another compared to non-significant paralogs (Fig. 2O-P). Considered individually, significant paralog vulnerabilities were members of smaller gene families, exhibited stronger evolutionary conservation, and were more likely to localize to the same cellular compartment compared to non-significant paralogs (Fig. 2Q-S). Collectively, these features suggest that aneuploidy-associated paralog vulnerabilities represent a distinct class of conserved genes from small families that serve redundant cellular functions.

### Aneuploidy-associated paralog vulnerabilities exhibit minimal transcriptional compensation in response to chromosome loss

Beyond the differences in functional properties described above, we speculated that significant and non-significant paralogs could exhibit distinct responses to the loss of a chromosome. Aneuploid cancer cells can partially buffer chromosome loss by increasing expression from retained gene copies, although the mechanisms underlying this compensation remain unclear^3,33–35^. We therefore compared chromatin and transcriptional responses to chromosome loss between significant and non-significant paralog vulnerabilities.

We first analyzed ATAC-seq data from human cancers, comparing chromatin accessibility around transcription start sites (TSSs) for genes encoded on copy-neutral and lost chromosomes (Fig. 3A)^36^. Genes on lost chromosomes exhibited higher ATAC-seq signal at their TSSs, suggesting that increased promoter accessibility following chromosome loss could contribute to dosage compensation (Fig. 3B). However, this association differed between the two classes of paralogs. While non-significant paralogs exhibited a pronounced increase in accessibility in the copy-loss state, this increase was significantly attenuated for paralog vulnerabilities (Fig. 3C-D). Permutation testing confirmed that the difference in ATAC-seq signal between copy-neutral and copy-loss states was significantly smaller for the latter group (P < 0.0002). We next examined H3K27 acetylation using HiChIP as a marker of active chromatin^37^. Chromosome loss was associated with a broad increase in H3K27ac around TSSs, but this increase was likewise attenuated around paralog vulnerabilities (Fig. 3E-F; P < 0.04).

**Figure 3.**
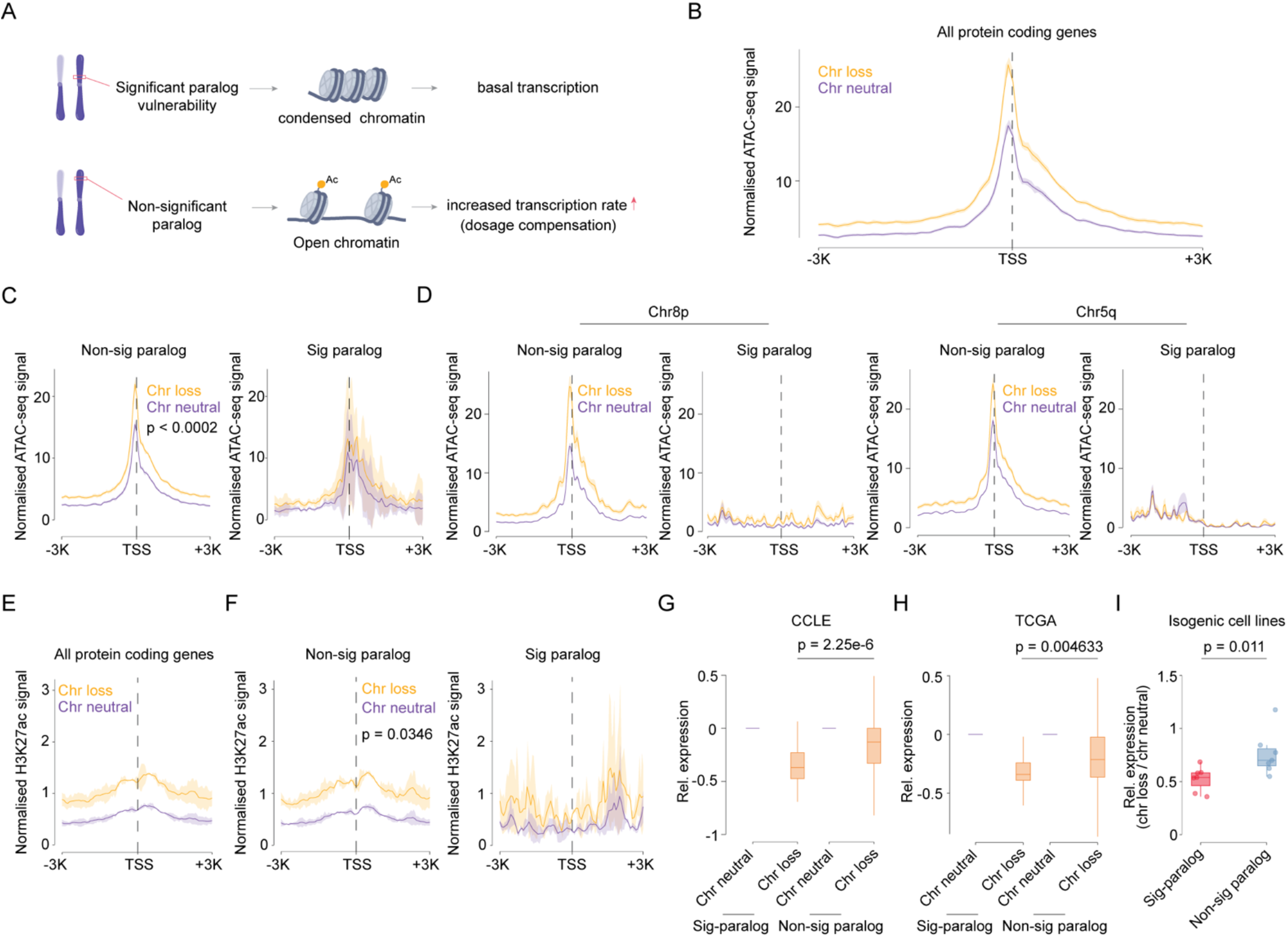
Chromatin and transcriptional responses to chromosome loss differ between paralogs with and without associated dependencies. (A) Proposed model of transcriptional compensation following chromosome loss. Limited chromatin accessibility constrains compensatory expression of paralogs whose loss creates a dependency on their partner, whereas increased accessibility permits upregulation of paralogs without an associated dependency. (B) Normalized ATAC-seq profiles around transcription start sites (TSSs) of protein coding genes when those genes are encoded on chromosomes that have been lost (orange) or on copy-neutral chromosomes (purple). (C) Normalized ATAC-seq profiles around transcription start sites (TSSs) of non-significant (left) and significant (right) paralogs. Profiles are shown for tumors with loss of the corresponding chromosome arm (orange) or neutral copy-number status (purple). (D) Normalized ATAC-seq profiles around the TSSs of paralogs on chromosome arms 8p and 5q, stratified by significance and chromosome copy-number status. (E) Normalized H3K27ac profiles around transcription start sites (TSSs) of protein coding genes when those genes are encoded on chromosomes that have been lost (orange) or on copy-neutral chromosomes (purple). (F) Normalized H3K27ac profiles around transcription start sites (TSSs) of non-significant (left) and significant (right) paralogs. Profiles are shown for tumors with loss of the corresponding chromosome arm (orange) or neutral copy-number status (purple). (G) Relative expression of significant and non-significant paralogs in cell lines with chromosome loss, normalized to the corresponding chromosome-neutral groups. The indicated P value compares relative expression between the two chromosome-loss groups. (H) Relative expression of significant and non-significant paralogs in patient tumors with chromosome loss, normalized to the corresponding chromosome-neutral groups. The indicated P value compares relative expression between the two chromosome-loss groups. (I) Relative expression of significant and non-significant paralogs in engineered isogenic cell lines that had lost individual chromosome arms, normalized to their copy-neutral control cell lines.

Finally, we directly compared paralog gene expression between copy-neutral and chromosome-loss states. In both cancer cell lines (CCLE) and in patient tumors (TCGA), paralog vulnerabilities exhibited a greater reduction in expression following chromosome loss compared to non-significant paralog genes (Fig. 3G-H). We extended this analysis to published RNA-seq data from engineered cell lines harboring losses of individual chromosome arms and their isogenic copy-neutral controls. As we observed with genetically-distinct tumors and cell lines, chromosome loss resulted in a significantly greater decrease in the expression of significant paralog genes compared to non-significant paralog genes (Fig. 3I; P < 0.02). Together, these findings suggest that cells compensate for chromosome loss by increasing chromatin accessibility and transcription from the remaining chromosome. However, this response is muted for certain paralogs, which creates a selective dependency following chromosome loss for their corresponding paralogs that are encoded elsewhere in the genome.

### Chromosome loss events establish CYCLOPS-like vulnerabilities

We noted that 28 genetic vulnerabilities associated with aneuploidy were genes directly encoded on the lost chromosome (Fig. 4A and Table S1). We hypothesized that these represented CYCLOPS genes, in which the loss of a chromosome decreased the dosage of an essential gene and rendered cells highly-sensitive to the ablation of that gene’s remaining copy^30^. For instance, cell lines harboring a deletion of chromosome 8p exhibited a highly-significant increase in dependency scores for *POLR3D*, a subunit of RNA polymerase encoded on that chromosome, compared to cell lines that were neutral for chromosome 8p (Fig. 4B-E). CYCLOPS genes were enriched for diverse biological processes, including protein glycosylation (*DPAGT1*, *ALG1*) and splicing (*RBM42*, *SNRPB*)(Fig. 4F and Table S2B). Unlike the paralog vulnerabilities, CYCLOPS genes typically lacked any closely-related family members in the genome: 64% of CYCLOPS genes were singletons compared to 34% of other genes encoded on lost chromosomes (Fig. 4G; P < 0.002, F test). Correspondingly, CYCLOPS genes exhibited borderline-essential dependency scores, even in diploid cell lines, and the dependency scores of CYCLOPS genes were globally correlated with their expression levels (Fig. 4H-I). Finally, as we observed with the paralog vulnerabilities, CYCLOPS genes exhibited less dosage compensation when encoded on a lost chromosome compared to other genes (Fig. 4J). Together, these findings suggest that CYCLOPS-like vulnerabilities arise from insufficient compensation for the reduced dosage of essential genes following chromosome loss. The absence of closely related paralogs for these genes further limits the cell’s capacity to tolerate reduced gene function, leaving cells highly sensitive to suppression of the remaining allele.

**Figure 4.**
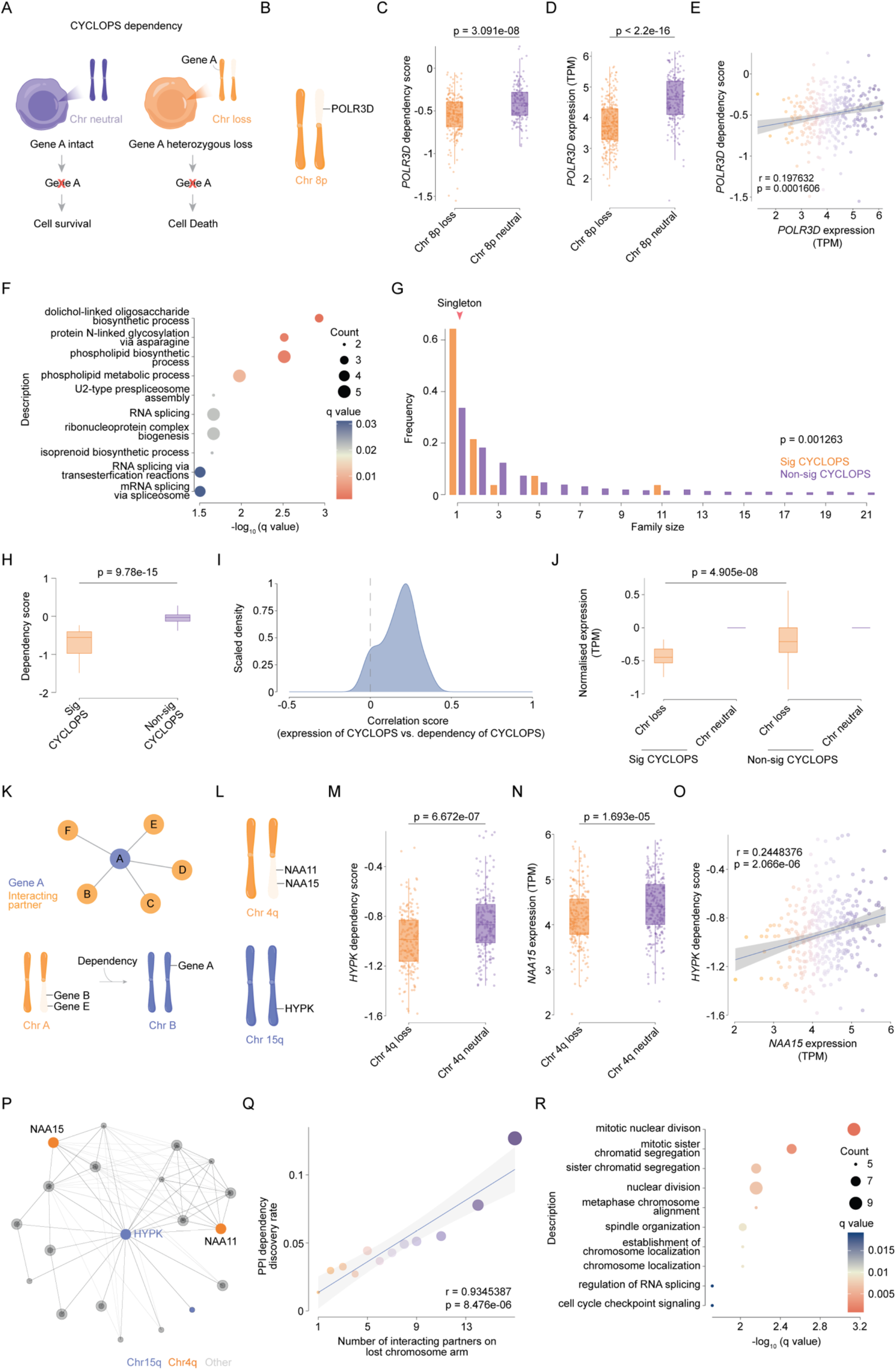
Chromosome loss events are associated with CYCLOPS-like and protein-interaction-associated dependencies. (A) Model of a CYCLOPS dependency. Reduced copy number of a gene following chromosome loss increases sensitivity to further suppression of that gene. (B) Location of *POLR3D* on chromosome 8p, illustrating its reduced copy number following 8p loss. (C) *POLR3D* dependency scores in cancer cell lines with chromosome 8p loss or neutral 8p copy-number status. (D) *POLR3D* expression in cancer cell lines with chromosome 8p loss or neutral 8p copy-number status. (E) Relationship between *POLR3D* expression and *POLR3D* dependency scores. Cell lines with chromosome 8p loss are shown in orange, and those with neutral 8p copy-number status are shown in purple. A fitted regression line is plotted against the data. (F) Gene ontology enrichment of significant CYCLOPS dependencies. Complete results are included in Table S2B. (G) Distribution of gene family sizes among genes identified as significant or non-significant CYCLOPS dependencies. (H) Dependency scores among genes identified as significant or non-significant CYCLOPS dependencies. (I) Distribution of correlation coefficients among genes identified as significant CYCLOPS dependencies. The dashed line indicates a correlation coefficient of zero. (J) Relative expression of significant and non-significant CYCLOPS genes in cell lines with chromosome loss, normalized to the corresponding chromosome-neutral groups. The indicated P value compares relative expression between the two chromosome-loss groups. (K) Model of an interactor dependency. Chromosome loss reduces the dosage of genes encoding interaction partners of a protein encoded elsewhere in the genome, increasing dependency on that protein. (L) Chromosomal locations of *NAA11* and *NAA15* on 4q and *HYPK* on 15q, illustrating the reduction in *NAA11* and *NAA15* copy number following 4q loss. (M) *HYPK* dependency scores in cancer cell lines with chromosome 4q loss or neutral 4q copy-number status. (N) *NAA15* expression in cancer cell lines with chromosome 4q loss or neutral 4q copy-number status. (O) Relationship between *NAA15* expression and *HYPK* dependency scores. Cell lines with chromosome 4q loss are shown in orange, and those with neutral 4q copy-number status are shown in purple. A fitted regression line is plotted against the data. (P) Protein interaction network containing HYPK and its interaction partners. Proteins encoded on chromosome 15q are shown in blue, those encoded on chromosome 4q are shown in orange, and other proteins encoded on other chromosomes are shown in grey. (Q) Relationship between the number of interaction partners encoded on a lost chromosome arm and the frequency of significant interactor dependencies. A fitted regression line is plotted against the data. (R) Gene ontology enrichment of significant protein-interaction dependencies. Complete results are included in Table S2C.

### Chromosome loss events create vulnerabilities by perturbing protein-protein interaction networks

We found that 72 genetic vulnerabilities associated with aneuploidy were genes that encoded proteins that interacted with proteins encoded on the lost chromosome, and in which there was a significant correlation between the expression level of the latter protein and the dependency score of the former protein (Fig. 4K and Table S1). For instance, the genes NAA11, NAA15, and HYPK encode components of the Nα-acetyltransferase A protein complex^38^. NAA11 and NAA15 are encoded on chromosome 4q, and loss of that chromosome arm is associated with a significant increase in the dependency of HYPK (Fig. 4L-P). As genes that encode interacting proteins are randomly scattered throughout the genome, we considered the possibility that these dependencies were unrelated to the interacting proteins highlighted by our analysis. However, if protein-protein interactions involving proteins encoded on a lost chromosome were the underlying cause of these vulnerabilities, then we hypothesized that proteins that interact with multiple proteins encoded on a lost chromosome would be more likely to be identified as aneuploidy-specific vulnerabilities. Consistent with this hypothesis, we observed a significant correlation between the number of proteins encoded on a lost chromosome that a protein interacted with and the likelihood that the gene encoding that protein was a selective dependency in those cells (Fig. 4Q; R = 0.93, P < 10^-5^). Gene ontology enrichment analysis revealed that these interaction-associated vulnerabilities were highly enriched for genes associated with chromosome segregation (*BUB3*, *ANAPC15*) and splicing (*RBM25*, *RBM22*)(Fig. 4R and Table S2C). These observations indicate that chromosome losses reshape protein–protein interaction networks, creating specific dependency network hubs that interact with multiple proteins encoded on the lost chromosome.

### Validation of *PPP2CA* as a vulnerability in cells with 8p-loss

In our CRISPR analysis, the most significant genetic vulnerability associated with an autosomal aneuploidy was the increased essentiality of the phosphatase *PPP2CA* in cancer cell lines that had lost a copy of chromosome 8p. We noted that chromosome 8p harbors a paralog of *PPP2CA* called *PPP2CB*, and low expression of *PPP2CB* was broadly correlated with increased sensitivity to *PPP2CA* ablation (Fig. 2D-G). To validate the biological significance of our analysis, we examined two sets of isogenic cell lines (AALE lung epithelial cells and HLF liver cancer cells) that were either disomic for chromosome 8p or in which a single copy of chromosome 8p had been eliminated using CRISPR (Fig. 5A)^29,39^. Read-depth analysis of low-pass whole-genome sequencing and TaqMan-based qPCR using a *PPP2CB* probe verified the 8p disomy in the parental cell lines and 8p monosomy in each derived clone (Fig. 5B-C and S4A-B). qRT-PCR demonstrated that the loss of 8p decreased the expression of *PPP2CB* by 2.9-fold in AALE cells and 1.8-fold in HLF cells, consistent with our data indicating that the paralog vulnerabilities identified by our analysis exhibit minimal transcriptional compensation (Fig. 5D and S4C). We also observed a moderate upregulation of PPP2CA in 8p-loss AALE cells but not in 8p-loss HLF cells (Fig. 5E and S4D).

**Figure 5.**
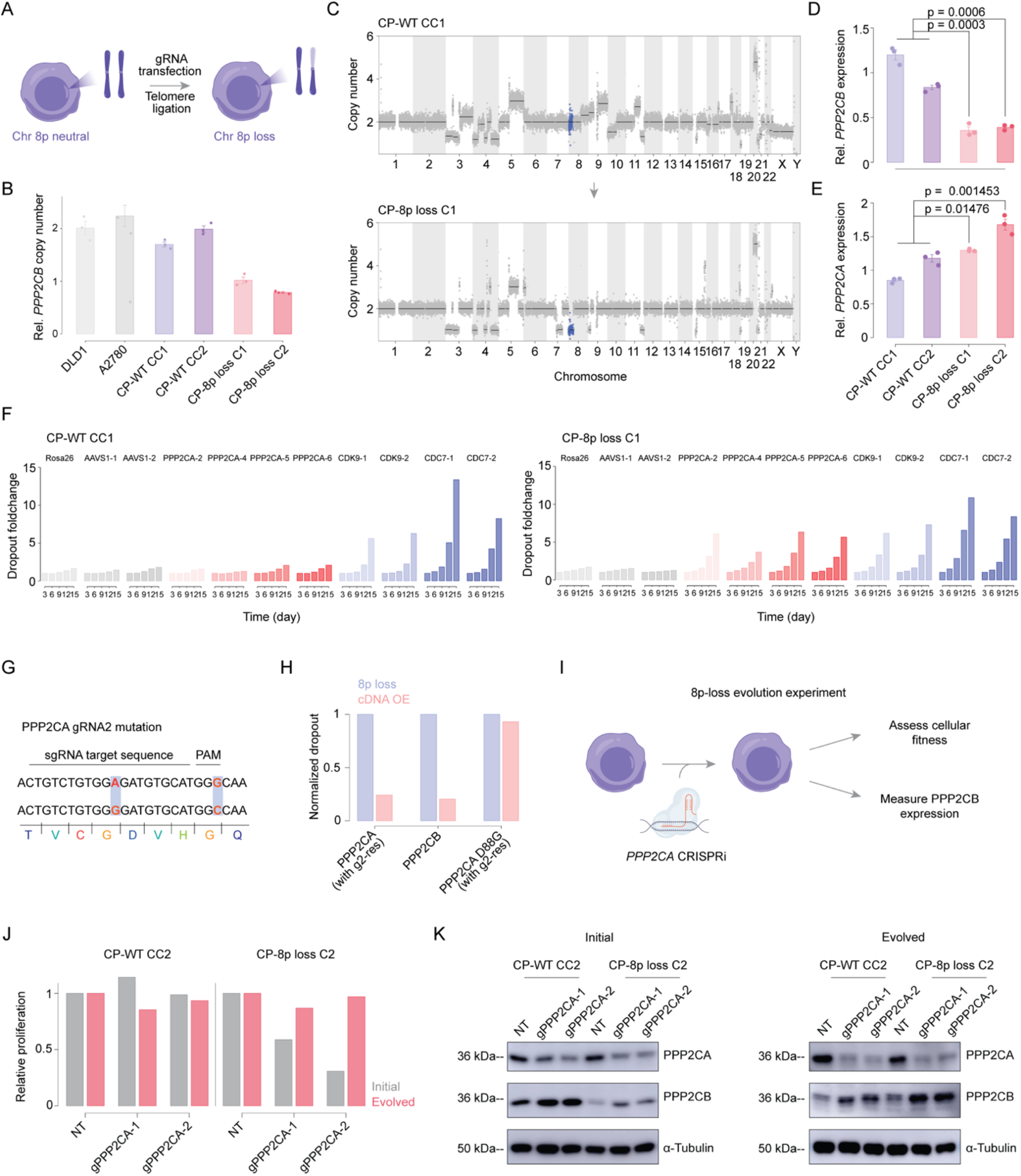
Chromosome 8p loss creates a selective dependency on PPP2CA. (A) Figure depicting the generation of isogenic cells with chromosome 8p loss using gRNA transfection and telomere ligation^9^. (B) TaqMan measurements of *PPP2CB* copy number in control 8p-neutral cancer cell lines (DLD1 and A2780), in 8p-neutral AALE cells (CP-WT CC1 and CP-WT CC2), and in 8p-loss AALE cells (CP-8p loss C1 and CP-8p loss C2). (C) Genome-wide copy-number profiles generated by whole-genome sequencing of CP-WT CC1 and CP-8p loss C1 cells. Chromosome 8p is highlighted in blue. (D) Relative *PPP2CB* expression in two CP control clones and two isogenic 8p-loss clones. (E) Relative *PPP2CA* expression in two CP control clones and two isogenic 8p-loss clones. (F) CRISPR competition assays in CP-WT CC1 and CP-8p loss C1 cells over 15 days following targeting with four independent *PPP2CA* gRNAs. *Rosa26* and *AAVS1* guides served as negative controls, and *CDK9* and *CDC7* guides served as pan-essential positive controls. Complete results are presented in Table S3. (G) Synonymous substitutions introduced into the *PPP2CA* cDNA to confer resistance to PPP2CA gRNA 2. (H) Rescue assays measuring *PPP2CA*-targeting gRNA-induced dropout in CP-8p loss C2 cells with or without ectopic expression of wild-type *PPP2CA*, *PPP2CB* or phosphatase-dead *PPP2CA(D88G)*. Both PPP2CA constructs carry the guide-resistance substitutions shown in G. Dropout is normalized to the 8p-loss cells without the corresponding rescue construct. (I) Experimental design for assessing adaptation to sustained *PPP2CA* suppression by CRISPR interference (CRISPRi). (J) Relative proliferation of CP-WT CC2 and CP-8p loss C2 cells carrying a non-targeting guide (NT) or either of two *PPP2CA*-targeting CRISPRi guides, measured at the initial (grey) and evolved (pink) time points. (K) Immunoblots of PPP2CA and PPP2CB in the cells shown in J at the initial and evolved time points. α-tubulin served as a loading control.

To examine the relationship between 8p copy number and *PPP2CA* essentiality, we performed CRISPR competition assays, in which we transduced 8p-disomic and 8p-monosomic cells with GFP-expressing gRNA vectors and then monitored the loss of fluorescent signal over time^40^. As expected, gRNAs targeting the non-coding loci *AAVS1* and *Rosa26* exhibited minimal depletion over five passages in culture, while gRNAs targeting the pan-essential cell cycle genes *CDC7* and *CDK9* were rapidly lost from all cell populations (Fig. 5F, S4E, and Table S3). Strikingly, we observed a significant difference in the response to *PPP2CA*-targeting gRNAs based on the copy number of chromosome 8p: in both 8p-disomic cell lines, gRNAs targeting *PPP2CA* were tolerated with no apparent decrease in fitness, while the same gRNAs were significantly depleted in both 8p-monosomic clones, indicating significant lethality (Table S3). To verify that these results were an on-target effect of *PPP2CA* ablation, we transduced an 8p-monosomic AALE clone with gRNA-resistant *PPP2CA* cDNA, and we found that this construct rescued the lethality of the *PPP2CA*-targeting gRNA (Fig. 5G-H). Viability required PPP2CA’s phosphatase activity, as a construct encoding the catalytically-inactive PPP2CA^D88G^ allele was unable to rescue the effects of *PPP2CA* ablation (Fig. 5H)^41^. Finally, transduction with wild-type *PPP2CB* cDNA also restored viability in 8p-monosomic cells, demonstrating that the selective dependency on *PPP2CA* is a direct consequence of decreased *PPP2CB* dosage and that the two paralogs perform at least partially interchangeable functions.

### Adaptation following *PPP2CA* suppression in cells with 8p-loss

While our competition experiments demonstrated that the loss of chromosome 8p establishes a selective genetic dependency on *PPP2CA*, these assays specifically measured the acute response to complete *PPP2CA* ablation. We noted that 8p-loss cells retain a single functional copy of *PPP2CB*, which we hypothesized could serve as a genetic reservoir for adaptation to prolonged PPP2CA inhibition. We therefore sought to model how 8p-loss cells evolve during sustained suppression of *PPP2CA*.

We initially sought to establish stable cell lines following transduction with the PPP2CA-targeting gRNAs used in the competition assays above. However, we found that transducing 8p-loss cells with PPP2CA-targeting gRNAs was lethal, and we were unable to establish viable cell lines following selection for gRNA-expressing cells. We therefore opted to use CRISPR interference (CRISPRi) to suppress the expression of *PPP2CA* without totally ablating it (Fig. 5I)^42^. We transduced cells with a dCas9-KRAB construct along with either a non-targeting gRNA or two gRNAs targeting near the *PPP2CA* start site. The *PPP2CA*-targeting gRNAs resulted in an ∼80% decrease in the expression of *PPP2CA* mRNA (Fig. S4F). Next, we conducted two proliferation assays: one immediately following selection for gRNA-expressing cells and one after allowing those cells to grow for 18 days. At the initial timepoint, PPP2CA suppression resulted in a ∼50% decrease in cell proliferation in 8p-loss cells, while proliferation of the 8p-neutral cells was unaffected. However, by day 18, the 8p-loss cells had recovered, and there was no significant difference in proliferation between the gNT-transduced and the gPPP2CA-transduced cell populations (Fig. 5J).

We suspected that this adaptation could be caused by an increase in *PPP2CB* expression following prolonged PPP2CA suppression. Interestingly, qPCR analysis revealed that *PPP2CB* mRNA levels remained largely unchanged in the evolved 8p-loss cell populations (Fig. S4G). However, western blotting demonstrated a progressive increase in PPP2CB protein levels in the 8p-loss cells. At 20 days following selection for the gRNA-transduced populations, PPP2CA protein remained suppressed in both wild-type and 8p-loss cell lines, but PPP2CB protein expression appeared significantly upregulated, and wild-type and 8p-loss cell lines expressed comparable levels of PPP2CB (Fig. 5K). These results suggest that 8p-loss cells can adapt to sustained *PPP2CA* suppression by restoring PPP2CB protein abundance through a post-transcriptional mechanism. Thus, although reduced *PPP2CB* dosage creates an initial dependency on *PPP2CA*, the residual *PPP2CB* allele provides a potential route to adaptation that can blunt the long-term effects of partial PPP2CA inhibition.

### Loss of the Y chromosome is a robust biomarker for *DDX3X* dependency

Our experiments with PPP2CA verified that chromosome loss events can establish specific paralog vulnerabilities, but they also raised the possibility that compensatory upregulation of the monosomic paralog could blunt its long-term efficacy as a therapeutic strategy. We therefore turned our attention to a class of genomic alterations in which upregulation of the lost paralog is not physically possible: loss of the Y chromosome. As LOY results in a nullizygous deletion of the genes encoded on the Y chromosome, any resulting paralog dependency cannot be overcome by re-expression of the lost gene. Indeed, the two most significant vulnerabilities identified by our analysis were *EIF1AX* and *DDX3X*, which are X chromosome-encoded paralogs of Y chromosome genes^43^. *EIF1AX* encodes a translation initiation factor that lacks a druggable enzymatic domain, and the protein that it encodes differs from its Y chromosome paralog *EIF1AY* by only a single amino acid substitution. Thus, developing a therapeutic strategy to selectively inhibit EIF1AX while sparing EIF1AY could represent an intractable pharmacological challenge. In contrast, *DDX3X* encodes an ATP-dependent RNA helicase, and helicases have attracted growing attention in recent years as promising targets for cancer therapy^44,45^. Moreover, DDX3X differs from its Y chromosome-encoded paralog DDX3Y along 8% of its sequence, potentially facilitating the development of paralog-specific inhibitors. We therefore decided to focus our efforts on characterizing *DDX3X* as a specific vulnerability in LOY cancers.

As a first step toward investigating DDX3X as a cancer vulnerability, we analyzed DepMap screening data to determine whether LOY is the best available biomarker for predicting *DDX3X* dependency, or whether other genomic or molecular features could enhance our ability to identify *DDX3X*-dependent cancers (Fig. S5A). We examined the mutation, copy number, and transcriptional profiles of DepMap cell lines and compared them to each line’s *DDX3X* dependency. The biomarkers most strongly correlated with *DDX3X* dependency were decreased copy number and low expression of genes encoded on the Y chromosome, including *USP9Y*, *UTY*, and *DDX3Y* (Fig. S5B-D). Consistent with this result, an independent elastic net regression analysis identified LOY as the most significant biomarker associated with *DDX3X* dependency (Fig. S5E). The relationship between *DDX3X* dependency and LOY was also consistent across cancer lineages, and no single cancer type exhibited a disproportionate reliance on *DDX3X* (Fig. S5F). We did not observe a correlation between MSI status or mutational burden and *DDX3X* dependency (Fig. S5G-I). WGD-positive and highly aneuploid cancers did exhibit a greater dependence on *DDX3X*, but closer analysis revealed that both WGD and aneuploidy were themselves correlated with LOY status (Fig. S5J-M). Taken together, these analyses indicate that *DDX3X* dependency is a prominent vulnerability in LOY cancers across lineages, and that incorporating additional genomic or molecular features does not substantially improve upon LOY for identifying *DDX3X*-dependent cancers.

We next asked how the DDX3X-LOY vulnerability compared to other reported aneuploidy-associated dependencies. We identified 22 aneuploidy-associated genetic vulnerabilities that had been published in the literature. These genes included *KIF18A*, *UBR4*, *UBE2H*, and *BUB1B*, which have been identified as vulnerabilities in WGD-positive and/or highly-aneuploid tumors, as well as *NDE1*, *VPS4A*, *PKMYT1*, and *MAGOHB*, which have been identified as vulnerabilities in cancers harboring specific chromosomal alterations. We conducted a uniform analysis of each of these genes in the presence or absence of their published biomarker using the DepMap CRISPR screening dataset (Fig. 6A-B and Table S4). We found that 11 of these genes exhibited significantly greater dependency scores in the presence of their published biomarker. However, the magnitude and significance of the association between *DDX3X* dependency and LOY were greater than any of the 21 published hits (Table S4). These results suggest that, within the limitations of the DepMap dataset, the *DDX3X*-LOY connection is the strongest aneuploidy-associated vulnerability that has yet been reported.

**Figure 6.**
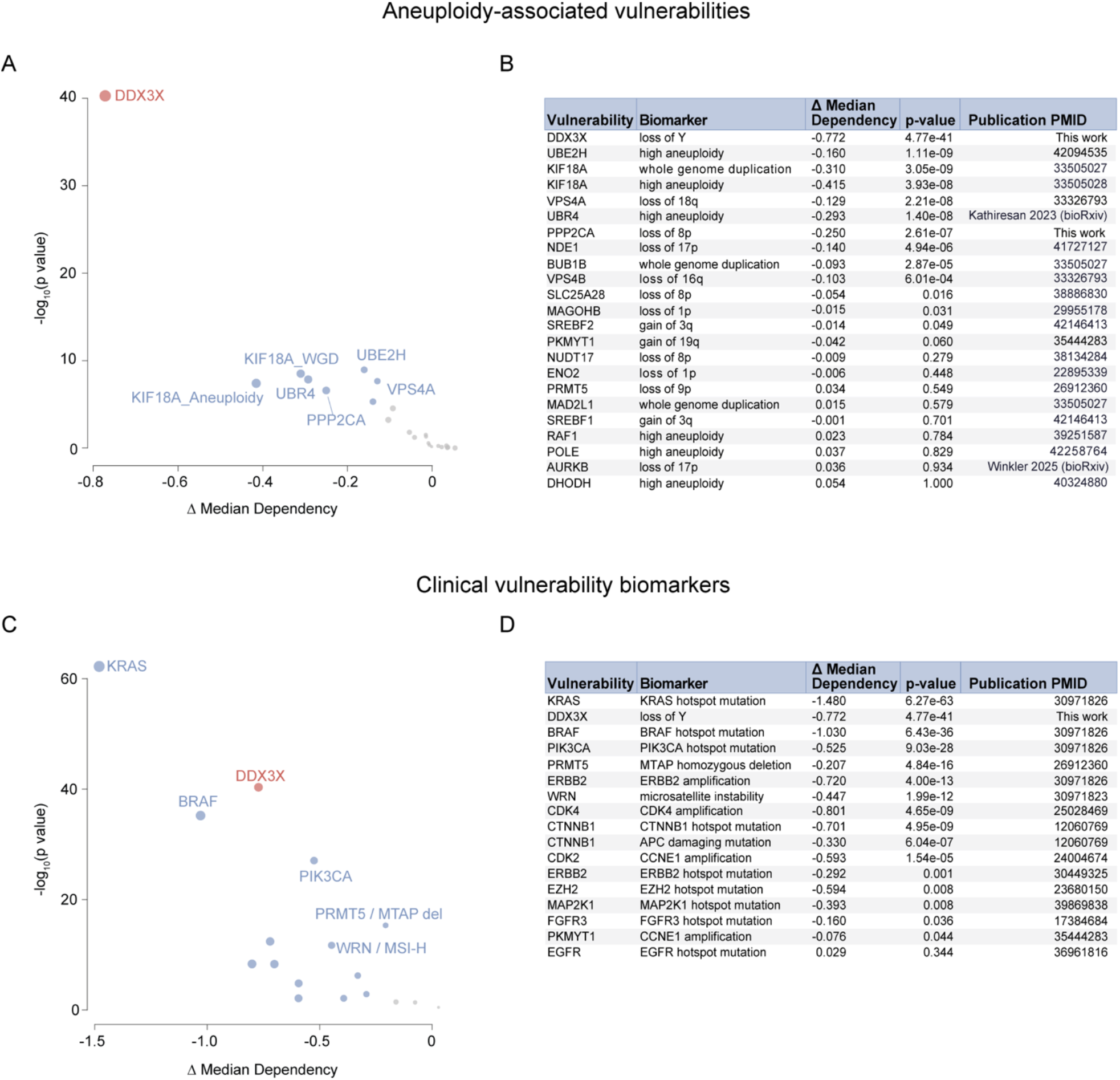
Comparing the DDX3X-LOY relationship with other aneuploidy-associated and clinical vulnerability biomarkers. (A) Comparison of the effect sizes and statistical significance of previously reported aneuploidy-associated dependencies with the DDX3X and PPP2CA dependencies described in this study. Each point represents a target–aneuploidy association. The x-axis shows the difference in median dependency scores between cell lines with and without the indicated aneuploidy feature; negative values indicate greater dependency in the biomarker-positive group. (B) A table summarizing the results shown in (A). (C) Comparison of the DDX3X–LOY association with established biomarker-associated dependencies. Each point represents a target–biomarker association, plotted by the difference in median dependency scores between biomarker-positive and biomarker-negative cell lines. (D) A table summarizing the results shown in (C).

Next, we evaluated *DDX3X* in comparison to vulnerabilities linked to other classes of genomic alterations, including several that have been established as clinically validated biomarkers. We found that the magnitude and significance of the *DDX3X*-LOY association were greater than many sensitivity biomarkers that are routinely used in clinical practice, including *PIK3CA* mutations as a biomarker for *PIK3CA* dependency, *ERBB2* amplifications as a biomarker for *ERBB2* dependency, and *EGFR* mutations as a biomarker for *EGFR* dependency (Fig. 6C-D and Table S4)^46^. *DDX3X*-LOY also exhibited a stronger association than several biomarkers that were discovered via functional genomic screening and have since advanced into clinical testing, including MSI status as a biomarker for *WRN* dependency and *MTAP* deletions as a biomarker for *PRMT5* dependency^47–50^. Although *DDX3X*-LOY was less significant than *KRAS* mutations as a biomarker for *KRAS* dependency, it was comparable to the use of *BRAF* mutations as a biomarker for *BRAF* dependency. Thus, when benchmarked against both clinically established and emerging genomic biomarkers, the magnitude of the *DDX3X*-LOY association supports further evaluation of DDX3X as a target in LOY cancers.

### Validating *DDX3X* as a selective dependency in LOY cancers

To validate the results of the CRISPR screening analysis, we applied our previously-described chromosome engineering pipeline to eliminate the Y chromosome from XY cancer cell lines^51^. We successfully established LOY (XO) clones from five genetically-distinct cancer cell lines derived from men: 786-O (renal adenocarcinoma), A2058 (melanoma), NALM6 (acute lymphoblastic leukemia), SNU1 (gastric adenocarcinoma), and THP1 (acute monocytic leukemia)(Fig. 7A)^51^. Read-depth analysis of low-pass whole-genome sequencing as well as a TaqMan-based qPCR using a *DDX3Y* probe verified loss of the Y chromosome from each line (Fig. 7B-C and S6A). qRT-PCR confirmed that *DDX3Y* expression was undetectable in each LOY clone, while the effects of LOY on *DDX3X* expression were variable across the cell lines (Fig. 7D-E).

**Figure 7.**
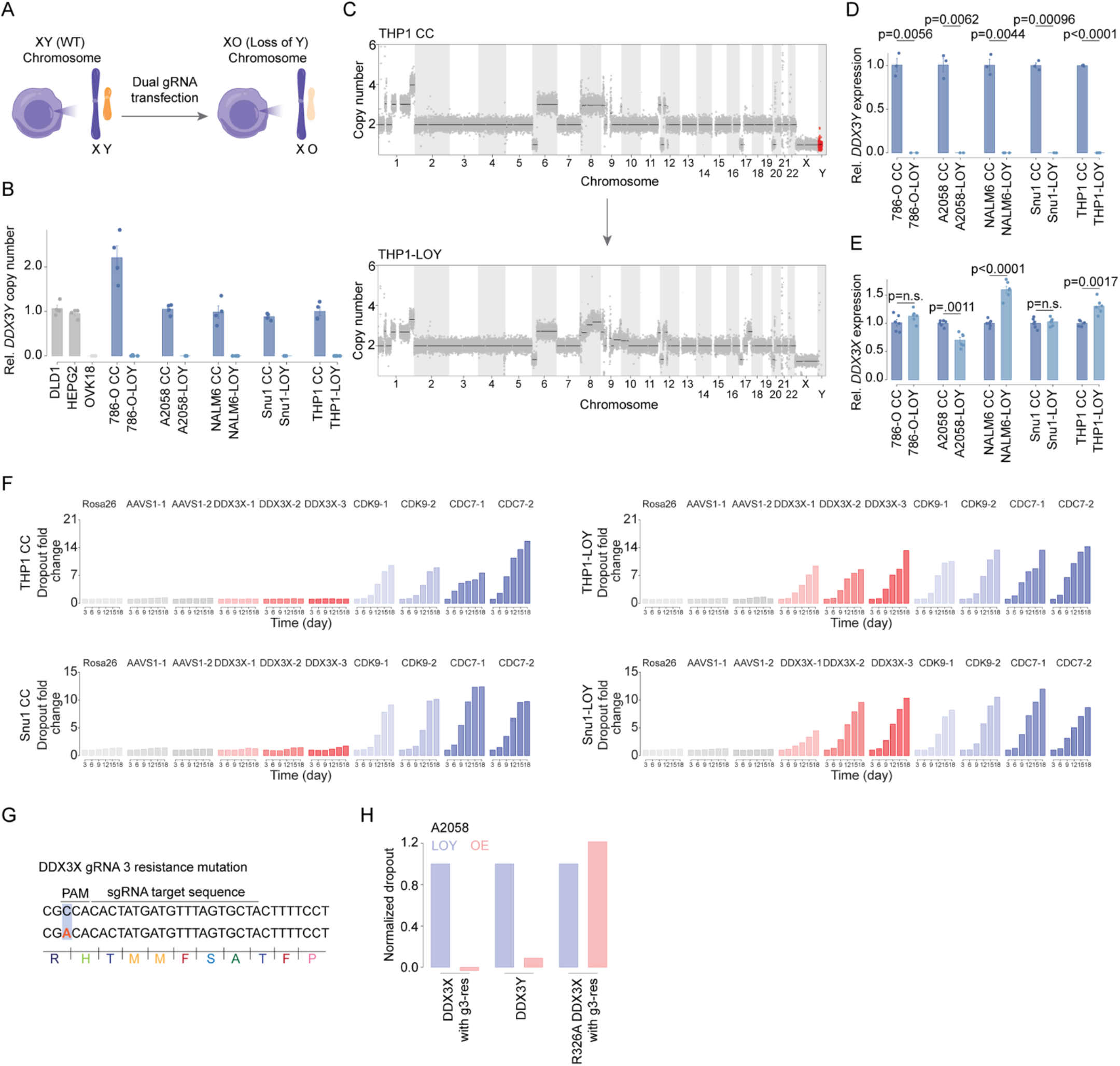
Loss of the Y chromosome creates a selective dependency on DDX3X. (A) Workflow for generating isogenic cells with Y chromosome loss using CRISPR gRNAs. (B) TaqMan measurements of *DDX3Y* copy number in control XY cancer cell lines (DLD1 and HEPG2), a control XX cancer cell line (OVK18), and five isogenic pairs of control (CC) and loss-of-Y (LOY) clones derived from 786-O, A2058, NALM6, Snu1 and THP1 cells. (C) Genome-wide copy-number profiles of THP1 CC and THP1-LOY cells generated by whole-genome sequencing. The Y chromosome is highlighted in red. (D) Relative *DDX3Y* expression in the five isogenic CC–LOY clone pairs. (E) Relative *DDX3X* expression in the same clone pairs. (F) CRISPR competition assays in isogenic THP1 and Snu1 CC and LOY cells over 18 days following targeting with three independent *DDX3X* gRNAs. *Rosa26* and *AAVS1* guides served as negative controls, and *CDK9* and *CDC7* guides served as pan-essential positive controls. (G) Synonymous substitution introduced into the *DDX3X* cDNA to confer resistance to DDX3X gRNA 3. (H) Rescue assays measuring *DDX3X*-targeting gRNA-induced dropout in A2058 LOY cells with or without ectopic expression of wild-type *DDX3X*, *DDX3Y* or helicase-dead *DDX3X(R326A).* Both DDX3X constructs carry the guide-resistance substitution shown in G. Dropout is normalized to LOY cells without the corresponding rescue construct.

Next, we performed CRISPR competition assays in five XY and five LOY clones using GFP-expressing gRNA vectors, as described above for *PPP2CA*. In all 10 clones, gRNAs targeting the negative control loci produced only modest changes in GFP-positive cells, whereas gRNAs directed against the pan-essential genes demonstrated a pronounced dropout (Fig. 7F, S6B, and Table S3). Transducing cells with DDX3X-targeting gRNAs resulted in significantly different outcomes based on the status of the Y chromosome: in each XY cell line, gRNAs targeting *DDX3X* were well-tolerated with no apparent fitness defect, whereas the same *DDX3X*-targeting gRNAs were strongly depleted in all five LOY clones, indicating significant lethality. Expression of gRNA-resistant wild-type *DDX3X* cDNA, but not cDNA encoding a helicase-dead allele of *DDX3X*, rescued the lethality of a DDX3X-targeting gRNA (Fig. 7G-H and Table S3). Expression of *DDX3Y* cDNA also restored viability in XO cells transduced with a DDX3X-targeting gRNA (Fig. 7H). Together, these experiments validate the DepMap analysis and establish *DDX3X* as a selective vulnerability across cancer types that arises from the functional redundancy between *DDX3X* and *DDX3Y*.

### Acute ablation of DDX3X triggers apoptosis and tumor regressions in mice

We next asked whether LOY cells could adapt to sustained *DDX3X* suppression, as we observed with *PPP2CA* in 8p-loss cells. However, both CRISPR and CRISPRi gRNAs targeting *DDX3X* were uniformly lethal in LOY cells, further underscoring the exquisite dependence of these cells on *DDX3X* function. As an orthogonal approach to achieve controlled depletion of DDX3X, we generated SNU1 LOY clones in which DDX3X was fused to the degron tag FKBP12^F36V^ (hereafter referred to as DDX3X-dTAG cells)(Fig. 8A)^52^. Treatment with either dTAG-13, which recruits cereblon, or dTAGV-1, which recruits VHL, caused a concentration-dependent loss of viability in DDX3X-dTAG cells (Fig. 8B). dTAGV-1 was more potent than dTAG-13 in these assays (IC50 of 2 nM versus 19 nM), and we therefore used dTAGV-1 for subsequent experiments.

**Figure 8.**
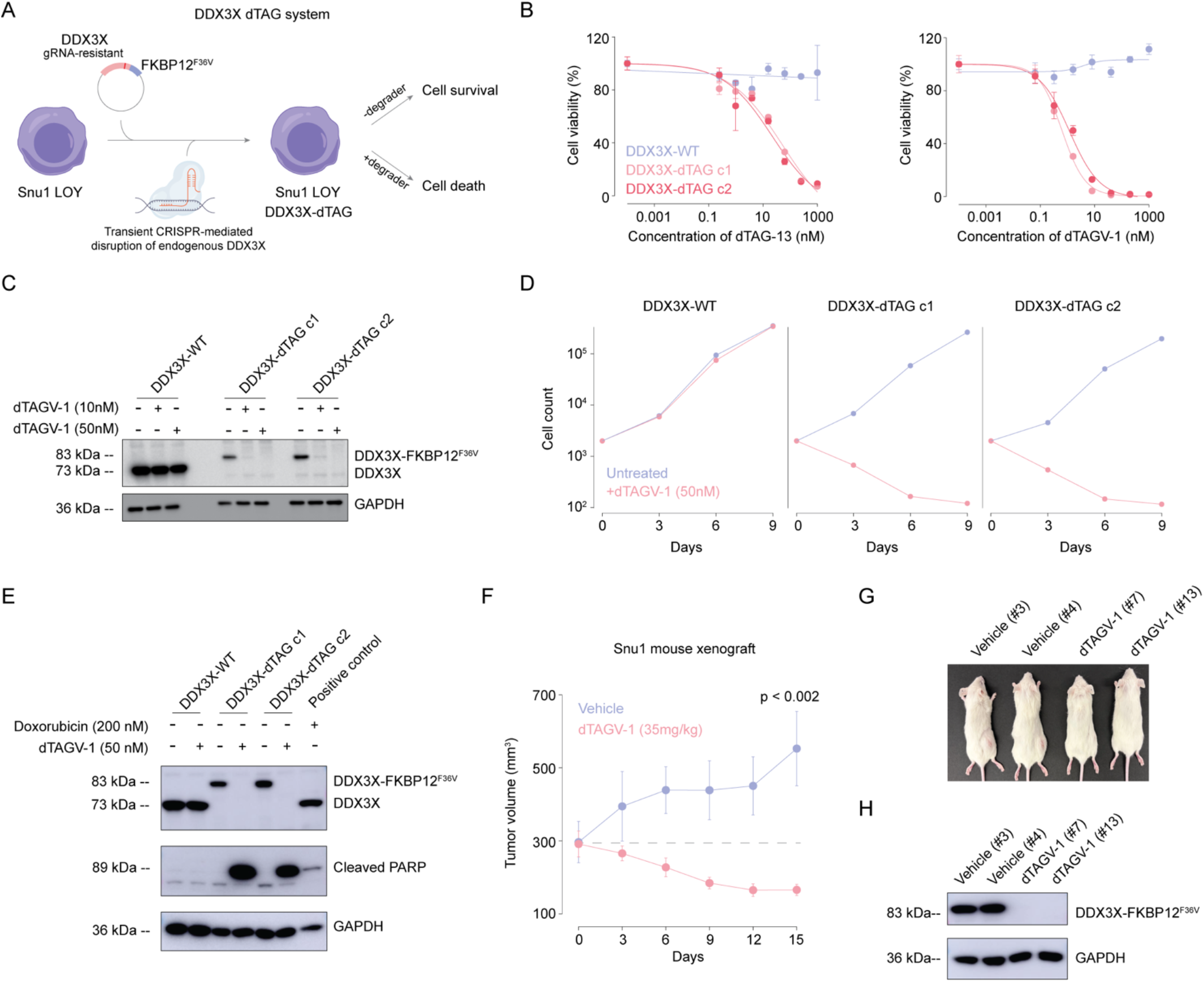
DDX3X ablation in LOY cancer cells triggers apoptosis and tumor regression. (A) Generation of DDX3X-dTAG derivatives of Snu1 LOY cells by expressing a gRNA-resistant DDX3X–FKBP12^F36V^ fusion and disrupting endogenous *DDX3X* using CRISPR. Addition of a dTAG degrader causes the acute depletion of the fusion protein. (B) Viability of parental DDX3X-WT and DDX3X-dTAG clones exposed to dTAG-13 (left) and dTAGV-1 (right). (C) Western blotting of DDX3X in DDX3X-WT and DDX3X-dTAG cells treated with 10 or 50 nM dTAGV-1. GAPDH serves as a loading control. (D) Cell counts in parental DDX3X-WT cells and DDX3X-dTAG clones treated with 50 nM dTAGV-1. (E) Western blotting of DDX3X and cleaved PARP in DDX3X-WT and DDX3X-dTAG cells treated with 50 nM dTAGV-1. Doxorubicin served as a positive control for PARP cleavage via apoptosis induction, and GAPDH served as a loading control. (F) Tumor volumes over 15 days in mice bearing Snu1 DDX3X-dTAG xenografts treated with vehicle or dTAGV-1 (35 mg/kg). The indicated P value compares the treatment groups on day 15. (G) Pictures of the mice from the xenograft experiment shown in (F). Cells were injected into the right side of each mouse. (H) Immunoblots of the DDX3X-dTAG fusion in xenograft samples from vehicle- and dTAGV-1-treated mice. GAPDH served as a loading control.

Western blotting confirmed that treatment of SNU1 LOY cells with dTAGV-1 completely depleted DDX3X protein (Fig. 8C). Next, we performed global proteomic profiling on DDX3X-dTAG cells, which revealed that DDX3X was the most strongly depleted protein after 24 hours of dTAGV-1 treatment (Fig. S7A-B and Table S6A). DDX3X ablation also decreased the abundance of chromosome segregation proteins including Aurora B, INCENP, and survivin, suggesting a role for DDX3X in controlling the expression of key mitotic regulators (Fig. S7C and Table S6B). Upregulated proteins included the tumor suppressors p53, CDKN2A (p14/p16), and CDKN1A (p21)(Table S6C).

We sought to determine whether DDX3X ablation causes cancer cell death. Over nine days of treatment, dTAGV-1 caused a progressive decline in the number of viable cells, indicating that acute DDX3X degradation causes a cytotoxic response (Fig. 8D). Consistent with this observation, dTAGV-1 treatment triggered cellular apoptosis, as assessed by western blotting for PARP cleavage (Fig. 8E). To determine whether DDX3X depletion could also affect established tumors, we generated subcutaneous DDX3X-dTAG xenografts in immunocompromised mice, and then we randomized the mice to receive either dTAGV-1 or vehicle. Treatment with dTAGV-1 triggered rapid tumor regressions, with tumor volume decreasing by an average of 43% over 15 days. In contrast, tumors in vehicle-treated mice increased in volume by an average of 86% over the same period (Fig. 8F-G). Western blotting of tumors collected from the treated mice revealed markedly reduced DDX3X protein levels, confirming target ablation (Fig. 8H). Together, these findings demonstrate that acute DDX3X ablation induces apoptotic cell death *in vitro* and regression of established tumors *in vivo*, providing proof-of-concept for targeting DDX3X in cancers with LOY.

## Discussion

Here, we establish that chromosome losses are a recurrent source of genetic vulnerabilities in cancer. By integrating chromosome copy number profiles with genome-scale CRISPR screens, we identified 389 dependencies associated with specific loss events and established that paralog insufficiency accounts for many of the strongest relationships. We validated two of these vulnerabilities in isogenic cells, demonstrating that loss of 8p creates a selective dependence on PPP2CA and that loss of the Y chromosome creates a selective dependence on DDX3X. The prevalence of chromosome losses in cancer and their relative scarcity in normal tissues make these alterations attractive biomarkers for potential therapies directed against aneuploid tumors.

Paralog redundancy provides a mechanism by which genes that are dispensable in normal tissue can become essential in cancer^53–55^. Previous research on this phenomenon has focused on instances in which one paralog is completely lost, either via homozygous deletion or via mutation-associated loss-of-heterozygosity. For instance, cancers harboring mutations or deletions of the chromatin remodeler *SMARCA4* become selectively dependent on the expression of its paralog, *SMARCA2*^56^. Similar functional redundancies have been reported between other paralogs in cancer, including *ENO1* and *ENO2*^57^, *STAG1* and *STAG2*^58^, and *ME2* and *ME3*^59^. Our identification of 37 aneuploidy-associated paralog vulnerabilities extends these observations and demonstrates that heterozygous deletions can also produce widespread vulnerabilities across diverse biological processes. Thus, complete elimination of a paralog is not required to compromise functional redundancy; reducing its dosage can be sufficient to increase dependence on its partner.

Moreover, paralogs whose reduced dosage was associated with a selective dependency on their partner exhibited attenuated increases in promoter accessibility and H3K27 acetylation, along with reduced transcriptional compensation following chromosome loss. These observations suggest that the capacity to increase transcription from the remaining allele helps determine the functional consequences of chromosome loss. When this response is insufficient, reduced expression can compromise functional redundancy, leaving cells dependent on a paralog encoded elsewhere in the genome. These results may explain why only a subset of paralog pairs gives rise to aneuploidy-associated vulnerabilities, despite chromosome loss events decreasing the copy number of hundreds of genes at once. Understanding why particular genes fail to mount an effective compensatory response could help identify additional therapeutic targets and clarify the regulatory mechanisms that allow cancer cells to tolerate aneuploidy.

We experimentally validated the *PPP2CA*-*PPP2CB* pair as one example of an aneuploidy-associated paralog vulnerability. These genes encode catalytic subunits of the PP2A complex, which reverses mitotic protein phosphorylation to coordinate chromosome segregation and cell cycle progression^60,61^. We demonstrated that the loss of chromosome 8p reduces PPP2CB expression and creates a selective requirement for PPP2CA phosphatase activity, while increasing PPP2CB expression rescues this vulnerability. However, the selective dependency on PPP2CA is transient: following prolonged suppression of PPP2CA, we found that 8p-loss cells eventually recovered and were able to proliferate at a wild-type rate. Interestingly, this recovery correlated with an increase in PPP2CB protein expression, while *PPP2CB* mRNA levels remained unchanged. Autoregulation of PP2A subunit expression via translational compensation has previously been documented^62^; whether this explains the upregulation of PPP2CB or occurs for other aneuploidy-associated paralog vulnerabilities is not currently known. Regardless of the mechanism, our experiments suggest a potential limit to the therapeutic benefits of targeting a heterozygous chromosome loss event, as the remaining allele can serve as a reservoir for cellular adaptation.

The most profound vulnerabilities in our analysis were associated with LOY. Previous studies have established the functional redundancy between several X- and Y-linked paralogs^43,63^, including DDX3X and DDX3Y^64–67^. Our findings demonstrate that this redundancy can be exploited to selectively target the LOY state across five genetically-distinct backgrounds. Beyond the strength of the underlying dependencies, LOY exhibits several desirable features as a potential therapeutic biomarker: loss of the Y chromosome is a binary event that results in the nullizygous deletion of the genes encoded on the Y chromosome. LOY is also irreversible, preventing tumors from adapting to therapeutic pressure via paralog re-expression. Clinically, LOY is straightforward to detect, either by copy number analysis of sequencing data or by FISH. Finally, LOY marks a patient population with substantial unmet medical need. Across tumor types, LOY is associated with aggressive disease, *TP53* mutations, and metastatic progression^10–12,14,68^, and LOY tumors establish an immunosuppressive microenvironment. These findings provide a strong biological rationale for targeting the LOY state in patients that currently lack effective treatment options.

One potential consideration for clinical translation is hematopoietic LOY, which has been observed in the leukocytes of aging men^69–71^. Hematopoietic LOY occurs in ∼7% of men aged 65 or younger, compared to 20% of men aged 86 years or older^72^. Patients with hematopoietic LOY may be more susceptible to treatment-induced toxicities, making this alteration a potential exclusion criterion for an LOY-targeting therapy. Alternatively, hematopoietic LOY has been associated with heart failure^73,74^, neurodegeneration^72,75,76^, and premature aging^70,77^, and transplant studies performed in mice have indicated that LOY cells directly cause increased mortality^78^. Thus, the selective elimination of LOY clones from the blood could potentially prove to be beneficial for patient health. Additional research will be required to determine how LOY-directed therapies affect normal hematopoiesis and whether the elimination of somatic LOY cells can alleviate the diseases associated with their accumulation.

Among potential vulnerabilities associated with LOY, we focused on DDX3X, as it encodes a druggable, ATP-dependent enzyme with 8% sequence divergence from its Y chromosome-encoded paralog^79,80^. DDX3X is a DEAD-box RNA helicase that promotes translation initiation, particularly for mRNAs with complex 5′ UTR structures^81,82^. It also regulates stress granule dynamics by resolving intermolecular RNA interactions that sequester mRNAs and limit their translation^83,84^. Within the limitations of the DepMap dataset, we found that the DDX3X–LOY association exhibits a greater effect size and statistical significance than 22 previously reported aneuploidy-associated vulnerabilities and was comparable to or stronger than many clinical biomarker–target relationships. Using an inducible degron, we further demonstrated that acute DDX3X depletion triggers apoptosis and regression of established LOY tumors, providing in vivo proof-of-concept for targeting this dependency.

Recent progress drugging helicases supports the feasibility of developing a pharmacological intervention targeting DDX3X. Two allosteric inhibitors of the DNA helicase WRN have recently advanced into testing in microsatellite-unstable cancers on the basis of a synthetic-lethal interaction first uncovered through CRISPR screening^47,48,85–87^. Inhibitors targeting several RNA helicases, including DHX8^88^, DHX9^89^, and eIF4A^90^, are also under active development. For DDX3X, a key challenge will be to develop potent compounds that spare DDX3Y, in order to minimize toxicity in healthy XY tissue. Ultimately, selective agents targeting DDX3X will be required to establish whether the genetic dependency documented in this work can be translated into a safe and effective therapy for LOY cancers.

## Methods

### Cell Culture

Human tumor cell lines were obtained from the ATCC or received as gifts from collaborating laboratories. The identity of all cell lines was verified by short tandem repeat (STR) profiling (University of Arizona Genetics Core). All cultures were routinely tested for *Mycoplasma* contamination using the MycoAlert® Mycoplasma Detection Kit (Lonza, cat. no. LT07-418) together with the MycoAlert® Assay Control Set (Lonza, cat. no. LT07-518).

The adherent A2058 melanoma cell line and HLF liver cancer cell line were maintained in DMEM (Gibco, cat. no. 11995065) supplemented with 2 mM L-glutamine (Gibco, cat. no. 25030149), 100 U/mL penicillin-streptomycin (Gibco, cat. no. 15140122), and 10% fetal bovine serum (FBS; Sigma-Aldrich, cat. no. 50-165-7327). Suspension cell lines THP1, Snu1, and NALM6, as well as the adherent 786-O renal carcinoma cell line, were cultured in RPMI-1640 medium (Gibco, cat. no. 11875093) supplemented with 2 mM L-glutamine, 100 U/mL penicillin-streptomycin, and 10% FBS. The AALE lung epithelial cell line (labeled as CP cell in the figures) was grown in SABM™ Basal Medium (Lonza, cat. no. CC-3118). All cells were maintained at 37 °C in a humidified atmosphere with 5% CO₂.

For cell counting, adherent cells were dissociated using TrypLE (Gibco, cat. no. 12-604-013), resuspended in growth medium, and pelleted at 300 × g for 5 min. Cell suspensions were mixed 1:1 with 0.4% trypan blue (Gibco, cat. no. 15250061) and viable cells were quantified using a Cellometer Auto T4 cell counter.

### Plasmid Constructs

#### CRISPR plasmid cloning

Guide RNAs for chromosome Y deletion were designed to target both the p-arm and q-arm regions using Benchling (www.benchling.com). sgRNAs targeting *DDX3X* and one set of sgRNAs targeting *CDK9* for competition assays were obtained from the Brunello library^91^. Additional sgRNAs targeting *CDC7* and *CDK9* were sourced from the Vakoc Human Kinase CRISPR library^92^. All sgRNAs were selected to target functional domains of the corresponding proteins. Negative control sgRNAs targeting *Rosa26* and *AAVS1* were obtained from the Sheltzer laboratory^93,94^, and a non-targeting sgRNA sequence was sourced from the Brunello library. sgRNAs targeting *PPP2CA*, *TYMS*, *FBXW11*, and *PPP2R1A* were obtained from DepMap to assess potential off-target paralog mutagenesis^95^. For competition assays, sgRNA sequences targeting *PPP2CA* were obtained from either Dharmacon or DepMap^95^. All sequences are provided in Table S5A.

Oligonucleotides were ordered from Integrated DNA Technologies (IDT), and sgRNAs for chromosome Y deletion and for competition assays were cloned into the Lenti-Cas9-gRNA-GFP vector (Addgene #124770) using BsmBI digestion (Thermo Scientific, cat. no. ER0451) as previously described^96^.

#### Cloning of CRISPRi plasmid

CRISPRi knockdown was performed using a two-plasmid system comprising pHR-UCOE-SFFV-Zim3-dCas9-P2A-Hygro (Addgene #188768) and pCRISPRia-v2 (Addgene #84832). Guide RNA sequences are listed in Table S5B. Guides were cloned into pCRISPRia-v2 according to the previously published protocol^97^.

#### Cloning of overexpression cDNA plasmid

The pLV_mCherry/Puro_EF1A_DDX3X (NM_001356.5) and pLV_mCherry/Puro_EF1A_DDX3Y (NM_004660.5) plasmids were obtained from VectorBuilder. Human *PPP2CA* cDNA (NM_002715.4) and *PPP2CB* cDNA (NM_001009552.2) were PCR-amplified from an MCF10A breast epithelial cell cDNA library. The *PPP2CA* and *PPP2CB* cDNAs were first cloned into pENTR1A (Addgene #17398) using SalI and NotI restriction sites (New England Biolabs, cat. no. R0138S and R0189S) and subsequently recombined into the pGen_EF1A_attR1_CmR_ccdB_attR2 destination vector (GenScript) using the Gateway cloning system according to the manufacturer’s instructions (Invitrogen, cat. no. 11791020). The plasmids and their Addgene numbers are provided in Table S5C.

#### cDNA Mutagenesis

Mutations to the g3 binding site in the *DDX3X* cDNA plasmid were created using the Q5 Site-Directed Mutagenesis Kit (NEB, cat. no. E0554S). The same kit was used to create an additional point mutation for the R326A helicase-dead *DDX3X* plasmid. Mutations to the g2 binding site in the *PPP2CA* cDNA plasmid were created using the Q5 Site-Directed Mutagenesis Kit (NEB, cat. no. E0554S). The same kit was used to create an additional point mutation for the D88G phosphatase-dead *PPP2CA* plasmid.

#### Cloning Degron Tagged DDX3X Overexpression Vector

A DDX3X isoform bearing sgRNA binding resistance mutations was obtained from the above modified pLV_mCherry/Puro_EF1A_DDX3X vector. Primers and experimental protocols for Gibson assembly and Hi-Fi assembly were designed by the NEBuilder Assembly Tool (nebuilder.neb.com). Gibson assembly was first used to assemble sgRNA-resistant DDX3X and FKBP12^F36V^ degron tag (Addgene #154287) into the pBABEpuro retroviral backbone (Addgene #40978). Hi-Fi assembly was then used to insert the combined DDX3X-FKBP12^F36V^ construct into Addgene #108100 in place of Cas9. All plasmids were validated by whole-plasmid sequencing (Plasmidsaurus).

#### Whole exome sequencing

Genomic DNA was extracted from cultured cells using the DNeasy Blood & Tissue Kit (Qiagen, cat. no. 69506) according to the manufacturer’s instructions. High-coverage whole exome sequencing (WES) was performed by Novogene. Libraries were prepared using standard Illumina-compatible workflows and sequenced to an average on-target depth of ∼50× per sample. Raw sequencing reads were quality-trimmed and filtered using fastp (v1.0.1) to remove adapter sequences and low-quality bases (minimum quality score threshold: -q 20)^98^. Filtered reads were aligned to the human reference genome hg38 using bwa-mem (v0.7.18) with default parameters. Resulting SAM files were converted to BAM format, sorted, and indexed using samtools (v1.20.0). Somatic variant calling was performed using the GATK (v4.6.1.0) best practices workflow^99^, with Mutect2 used for single nucleotide variant and indel detection^100^. Variants were filtered using recommended filtering parameters, removing all germline mutations using a panel of normals. Variant allele frequency (VAF) analyses were conducted for on-target and paralog off-target mutagenesis. For on-target mutagenesis, VAFs were calculated for all variants located within a ±10 bp window surrounding the CRISPR cut site. For potential paralog off-target mutagenesis, the mean VAF was calculated across all exonic variants mapping to the corresponding paralog gene. All downstream visualization and plotting were performed in R using custom scripts built with tidyverse (v2.0.0) package^101^.

### Chromosome loss clone generation

#### Loss-of-Y clones

Cells were transfected with gRNA CRISPR plasmids targeting the p and q arms of the Y chromosome near the centromere using Lipofectamine 3000 (Invitrogen, cat. no. L3000015). GFP+ cells were sorted on 96-well plates 48-72 hours after transfection. Clones were screened for Y-chromosome loss using TaqMan copy number probes.

#### 8p-loss clones

The 8p-loss AALE cells were previously described in ref^29^. The 8p-loss HLF cells were previously described in ref^39^.

#### TaqMan copy number assay

Genomic DNA was extracted and purified with the Qiagen DNeasy Blood & Tissue Kit (Qiagen, cat. no. 69506). TaqMan assays were conducted in quadruplicate using the TaqPath ProAmp Master Mix (Applied Biosystems, cat. no. A30867). The assay employed RNase P (Applied Biosystems, cat. no. 4403328) as the reference. Quantification was performed on the CFX OPUS 384 Real-Time PCR system (Bio-Rad). Copy number was normalized to the near-diploid cancer cell lines DLD1 and A2780. To detect Y chromosome loss, three Y chromosome probes and one chromosome 1q control gene were used. To detect chromosome 8p loss, one chromosome 8p probe was used. All probes are listed in Table S5D.

#### RT-qPCR

RNA was isolated and purified with the Qiagen RNeasy Mini Kit (Qiagen, cat. no. 74106). cDNA synthesis was carried out using Superscript IV VILO Master Mix (Invitrogen, cat. no. 11756500) or Transcriptor First Strand cDNA Synthesis Kit (Roche, cat. no. 04379012001). Quantitative PCR was conducted in triplicate for the target genes DDX3X, DDX3Y, PPP2CA and PPP2CB, as well as the control gene GAPDH, employing PowerTrack SYBR Green Master Mix for qPCR (Applied Biosystems, cat. no. A46109). The qPCR results were obtained using the CFX OPUS 384 Real-Time PCR system (Bio-Rad). Primers, designed via NCBI Primer-Blast, were ordered from IDT (Table S5E).

#### Karyotyping and copy number analysis

Genomic DNA was extracted from cultured cells using the DNeasy Blood & Tissue Kit (Qiagen, cat. no. 69506) following the manufacturer’s protocol. Low-coverage whole-genome sequencing (lcWGS) was performed by Azenta. Libraries were prepared using standard Illumina-compatible workflows. Raw sequencing reads were quality-trimmed and filtered using fastp (v1.0.1) to remove adapters, low-quality bases (-q 20)^98^. Filtered reads were aligned to the human reference genome hg38 using bwa-mem (v0.7.18) with default parameters. SAM files were converted to BAM format, sorted, and indexed using samtools (v1.20.0). Copy number profiles were inferred using CNVkit (v0.9.12), employing the *wgs* mode with circular binary segmentation (CBS) for segmentation^102^. Coverage normalization, bin-level log2 ratio estimation, and segmentation were performed according to the CNVkit documentation. All downstream visualization and plotting were performed in R using custom scripts built with tidyverse (v2.0.0) and DNAcopy (v1.80.0) packages.

#### Lentivirus Production and Cell Transduction

HEK293FT cells were transfected for lentivirus using the calcium-phosphate method^96^, as well as using Lipofectamine 3000 (Invitrogen cat. no. L3000015), or TRANS-IT (Mirusbio, cat. no. MIR 2300). Virus supernatant was harvested 48-72 hours post transfection using a 10 mL syringe and 0.45 µm filter (Millipore Sigma, cat. no. SLHVR33RS). Virus was aliquoted into 1.5 mL tubes and stored for future use or applied directly to cells with 4-10 μg/mL polybrene (Santa Cruz Biotechnology, Inc., cat no. SC-134220). The culture medium on target plates was changed 24 hours post-transduction.

#### GFP dropout competition assay

GFP competition assays were performed as previously described^40^. In brief, approximately 1.5 × 10⁵ cells were seeded in 12-well or 6-well plates and transduced with lentiviral vectors encoding Cas9, GFP, and a sgRNA targeting the gene of interest. Depending on the cell type, 0.5-1 mL of viral supernatant was added per well. Culture medium was replaced 24 h after transduction. At 72 h post-transduction, adherent cells were trypsinized, and a subset of cells was resuspended in 100 µL PBS for flow cytometry. Suspension cell lines were pelleted and similarly resuspended in 100 µL PBS. GFP⁺ cells were quantified on a CytoFLEX S flow cytometer (Beckman Coulter) using the B525 laser. Cells were replated after analysis, and the proportion of GFP+ cells was measured every 72 h for 5 or 6 time points (15- or 18-days total). Fold dropout was calculated as the ratio of the initial GFP+ fraction to the GFP+ fraction at each subsequent time point.

#### PPP2CA knockdown evolution assay

AALE cells with chromosome 8p loss or neutral chromosome 8p status, expressing either a non-targeting CRISPRi guide or a PPP2CA-targeting CRISPRi guide, were selected with puromycin for 5 days. Cells were seeded at 20,000 cells per well in six-well plates. Cell numbers were measured every 3 days using a CytoFLEX S flow cytometer and used to calculate proliferation rates over time. Proliferation rates were compared between chromosome 8p-loss and chromosome 8p-neutral cells for each guide condition. Cell pellets were collected at the start and end of the assay for quantitative PCR and immunoblot analysis of PPP2CA and PPP2CB RNA and protein expression, respectively.

#### Generation of DDX3X-Degron Clones

Cell lines were first stably transduced with the DDX3X-dTAG vector. Transduced populations were selected with puromycin to remove nontransduced clones. Transduced cells were then transiently transfected with GFP CRISPR plasmid number 3 to knock out endogenous DDX3X but not the exogenous DDX3X construct. 48 hours post transfection, cells were viability stained with 1 µM SytoxBlue (Molecular Probes #S34857) and incubated following reagent protocols. Stained cells were single-cell cloned via FACS, gating for GFP-positive and SytoxBlue-negative clones only.

#### Drug Curves

30,000 Snu1 cells were plated per well with 500 µL of RPMI media in a 24-well plate. dTAGV-1 (MedChemexpress #HY-145514D) or dTAG-13 (SelleckChem, cat. no. S9813) was added in triplicate to wells. Cell viability was assessed 96 hours post-treatment using a 1:1000 SytoxBlue stain (Fisher Scientific, cat. no. 50-113-7613) and the V450 laser of the CytoFLEX S flow cytometer (Beckman Coulter).

#### Western Blot

Cell pellets were lysed in RIPA buffer (Sigma-Aldrich, #R0278) supplemented with phosphatase inhibitor cocktail (Roche, cat. no. 49068450010) and protease inhibitor cocktail (Roche, cat. no. #04693116001). Protein concentrations were determined using the DC Protein Assay Kit (Bio-Rad, cat. no. #5000111). Equal amounts of protein were separated at 120 V on either 4-15% SDS-PAGE gels (Bio-Rad, cat. no. #4561086) or 10% Tris–glycine mini gels (Thermo Fisher Scientific, cat. no. #XP00105BOX). Proteins were transferred to PVDF membranes using either Trans-Blot Turbo transfer packs (Bio-Rad, cat. no. #1704156) with the Trans-Blot Turbo Transfer System or a Mini Trans-Blot cell (Bio-Rad). Membranes were blocked in 5% nonfat milk prepared in TBS containing 0.1% Tween-20 and incubated with primary antibodies overnight at 4 °C. After three 10-min washes in TBS-T, membranes were incubated with secondary antibodies for 1 h at room temperature. Membranes were washed again, incubated with Clarity Max Western ECL Substrate (Bio-Rad, #1705062) for 1 min, and imaged using an Amersham Imager 600 (Cytiva). The antibodies used are listed in Table S5F.

#### DDX3X degradation and proliferation assay

Snu1 LOY cells expressing degron-tagged DDX3X were seeded at 20,000 cells per well in six-well plates and treated with or without dTAGV-1. Treatment was renewed every 3 days. Cell numbers were quantified using a CytoFLEX S flow cytometer, and proliferation rates were calculated from cell counts collected over time.

### Quantitative global proteome profiling using LC/MS-MS

#### Sample preparation

Snu1-LOY and Snu1-LOY DDX3X-dTAG cells were treated for 24 hours with dTAGV-1 (50 nM) or DMSO in biological triplicate. Cells were harvested by scraping, washed twice with PBS and stored at -80°C until further use. Cell pellets were lysed thermally for 5 min at 95°C. After cooling at room temperature, freshly prepared lysis buffer (60 µL, 8 M urea, 150 mM NaCl, 100 mM HEPES pH 8.0 in MS-grade water) was added, and lysates were homogenized using probe ultrasonication (8 pulses, 25% amplitude). Total protein concentrations were quantified via Pierce BCA protein assay (Thermofisher, cat. no. 23225), and samples were normalized to 2 µg/µL (30 µg total) in lysis buffer. Samples were simultaneously reduced and alkylated by adding 10 µL of freshly prepared 10 mM Tris(2-carboxyethyl)phosphine hydrochloride (TCEP; Sigma-Aldrich, C4706) and 40 mM chloroacetamide (Sigma-Aldrich, C0267) and by shaking for 60 min at room temperature. Samples were then diluted by adding 120 µL CaCl_2_ (Sigma-Aldrich, C4901) in 20 mM HEPES, pH 8.0, to final concentrations of 0.9 M urea and 1 mM CaCl_2_ and digested overnight at 37°C (with shaking) using 1:100 (w/w ratio protease/lysate) Trypsin/LysC protease mix (Thermo, A40009). The digested samples were then acidified to pH ≤3 (confirmed by pH paper) by adding trifluoroacetic acid (TFA, final amount of 0.8%) and desalted using SOLAμ SPE HRP Peptide Desalting columns (Thermo 60209-001) with a Positive Pressure Manifold. Briefly, the desalting cartridges were activated with 200 μL of acetonitrile (ACN), then equilibrated twice with 200 μL wash buffer (2% ACN + 0.2% TFA in water). After loading peptides into the cartridges, samples were washed with 200 μL wash buffer (3 times) and eluted using 100 μL of 50% ACN with 0.1% formic acid (FA) in MS-grade water (twice). The desalted peptides were then immediately dried on a centrifugal vacuum concentrator (60°C, Thermo SPD120-115). The dried peptides were then reconstituted in 0.1% FA in water and analyzed by LC/MS-MS.

#### LC/MS-MS diaPASEF Acquisition

Reconstituted desalted peptides were analyzed using a nanoElute 2 UHPLC (Bruker Daltonics, Bremen, Germany) coupled to a timsTOF HT (Bruker Daltonics, Bremen, Germany) via a CaptiveSpray nano-electrospray source. The peptides were separated in the UHPLC using an Aurora Ultimate nanoflow UHPLC column with CSI fitting (25 cm x 75 µm ID, 1.7 µm C18; IonOptics AUR3-25075C18-CSI) at a flow rate of 400 nL/min with column temperature maintained at 50 °C using Mobile Phase A (MPA; 3% acetonitrile + 0.1% FA in water) and Mobile Phase B (MPB; 0.1% FA in ACN). The 70 min gradient started at 6% MPB while increasing to 17% MPB at 40 min, 25% MPB at 55 min, 34% MPB at 64 min, 85% MPB at 65 min, and finishing at 92% MPB at 70 min. The TIMS elution voltages were calibrated linearly with three points (Agilent ESI-L Tuning Mix Ions; 622, 922, 1,222 m/z) to determine the reduced ion mobility coefficients (1/K_0_). diaPASEF was performed using the MS settings 100 m/z for Scan Begin and 1700 m/z for Scan End in positive mode, the TIMS settings 0.70 V⋅s/cm^2^ for 1/K_0_ start, 1.30 V⋅s/cm^2^ for 1/K_0_ end, ramp time of 120.0 ms, 100% duty cycle, ramp rate of 7.93 Hz, and the capillary voltage set to 1600 V. diaPASEF windows from mass range 226.8 Da to 1226.8 Da and mobility range 0.70 1/K_0_ to 1.30 1/K_0_ were designed to provide 25 Da windows covering doubly and triply charged peptides as confirmed by DDA-PASEF scans, whereas singly charged peptides were excluded from the acquisition due to their position in the m/z-ion mobility plane.

#### Raw data processing

The raw diaPASEF files were processed using library-free analysis in FragPipe 24.0. DIA spectrum deconvolution was performed using diaTracer 2.2.1^103^ with the following default settings: (i) “Delta Apex IM” to 0.01, (ii) “Delta Apex RT” to 3, (iii) “RF max” to 500, (iv) “Corr threshold” to 0.3, and (v) mass defect filter enabled with offset set to 0.1. The reviewed UniProt Homo sapiens protein sequence database (01/06/2026; 20,453 entities) was supplemented with common contaminants and 50% decoys. In the MS Fragger 4.4.1 database search, the following settings were used: (i) initial precursor and fragment mass tolerances of 10 ppm and 20 ppm, respectively, (ii) enabled spectrum deisotoping, mass calibration, and parameter optimization, and (iii) isotope error set to “0/1/2”. For protein digestion, “stricttrypsin” for fully tryptic peptides was enabled with up to 1 missed cleavage, peptide length from 7 to 50, and peptide mass range from 500 to 5,000 Da. For modifications, methionine oxidation (2 max occurrences) and N-terminal acetylation (1 max occurrence) were set as variable modifications (maximum up to 3), while cysteine carbamidomethylation was set as a fixed modification. For validation, MSBooster (DIA-NN model) and Percolator were used for RT and MS/MS spectra prediction and PSM rescoring, while ProteinProphet (--maxppmdiff 2000000) was used for protein inference with FDR filtering (--picked --prot 0.01). The spectral library was then generated by FragPipe-SpecLib version 0.1.58 using default settings. Peptides were then quantified using DIA-NN 2.3.1 Academia with 1% FDR and QuantUMS high precision settings. The DIA-NN parquet report containing peptide quantification and scoring was then analyzed in R.

#### Statistical analysis

DIA-NN quantified peptides were then further filtered including (i) common contaminants and reverse sequences, (ii) 1% FDR filtering at the global.q.value (precursor Q value across all samples) and pg.q.value (protein group Q value in single injection), (iii) quantification of a given peptide in at least two replicates in one condition, and (iv) removal of singly charged peptides and non-proteotypic (unique) peptides. Protein intensities were then re-calculated using the MaxLFQ method^104^ provided in the DIA-NN R package. Differential statistics was then performed using the DEqMS R package^105^, which performs a LIMMA-moderated t-test with an adjustment for number of detected peptides per protein, to determine the p-value, fold change, and Benjamini-Hochberg adjusted p-value.

### Targeting DDX3X-dTAG in an in vivo tumor model

To assess the effects of DDX3X degradation on tumor growth *in vivo*, we established a xenograft tumor model with Snu1 DDX3X-dTAG c2. A total of 15x10^6^ cells were resuspended in a 1:1 ratio of Matrigel (Corning, 356237) and 0.9% saline and injected into the flank of female NOD.Cg-*Prkdc^scid^ Il2rg^tm1Wjl^*/SzJ (NSG) mice (Jackson Laboratory, cat. no. #005557). Mice were monitored for tumor formation, and once tumors became palpable, tumor dimensions were measured using digital calipers every three days. Tumor volume was calculated using the formula V= ½ x (length) x (width)^2^.

Once tumors reached an average volume of 250 mm^3^, mice were randomized into control and treatment groups based on tumor volume. Treatment with the dTAG degrader, dTAGV-1 TFA (TargetMol, #T40251) was then initiated. Tumor volume and body weight were monitored throughout the treatment period. dTAGV-1 TFA was formulated as previously described in Nabet et al., with the compound first dissolved in DMSO and then diluted with Solutol HS-15 (MedChemExpress, cat. no. #HY-Y1893) and then 0.9% saline with the final formulation containing 5% DMSO and 20% Solutol HS-15^52^. Mice received 35 mg/kg dTAGV-1 TFA or vehicle via intraperitoneal injections with twice-daily dosing until the end of the experiment. A minimum of 6 mice were used per group. All mouse protocols were approved by the Stanford Animal Care and Use Committee.

### Evaluation of off-target mutagenesis using DepMap constructs

To evaluate whether the gRNAs used in the DepMap result in off-target mutations in paralogous genes, guide RNA sequences from DepMap were cloned into lentiCRISPR v2-Blast (Addgene, #83480) by BsmBI digestion followed by Golden Gate assembly. We tested sgRNAs targeting FBXW11, PPP2CA, PPP2R1A and TYMS in A2780, HCT116 and MCF10A cells. A2780 cells were cultured in RPMI 1640, and HCT116 cells in DMEM. MCF10A cells were cultured in DMEM/F12 supplemented with cholera toxin (100 ng/ml; MilliporeSigma, C8052-0.5MG), epidermal growth factor (20 ng/ml; Gibco, AF-100-15-500UG), insulin (0.01 mg/ml; MilliporeSigma, I1882-100MG), hydrocortisone (500 ng/ml; MilliporeSigma, H0888-1G), horse serum (5%; Invitrogen, 16050-122), penicillin–streptomycin (1%) and transferrin (50 ng/ml, MilliporeSigma, T8158-100MG). Following transduction, cells were selected with blasticidin for 7 days at 5 µg/ml for MCF10A, 7 µg/ml for HCT116 and 4 µg/ml for A2780. Cells were then cultured in their respective growth media without blasticidin for a further 3 days before pellets were collected for whole-exome sequencing.

### Computational approaches

#### Chromosome loss dependency analysis

To assess chromosome loss-associated genetic dependencies in aneuploid cancers, we analyzed the genome-wide CRISPR loss-of-function screening dataset from DepMap, comprising 1,150 cancer cell lines (version 24Q2)^31^. Chromosome loss events for autosomes were obtained from ref^21^. For autosomal aneuploidies, we restricted our analyses to chromosome-arm loss events present in at least 150 cell lines to ensure adequate statistical power. To identify cancer cell lines with loss of the Y chromosome, we examined copy number data from CCLE, limiting our analysis to cell lines annotated as male. Loss of chromosome Y was defined as the absence of detectable copy number for at least 25 Y-linked genes.

Because aneuploid cancer cell lines harbor diverse combinations of chromosomal gains and losses, grouping cell lines solely by a specific chromosome-arm loss produced substantial imbalance in other chromosome-arm abnormalities between comparison groups. We recognized that such imbalances could potentially confound the identification of true chromosome-loss specific genetic dependencies. To mitigate this, we applied an aneuploidy score correction and restricted analyses to cell lines with an aneuploidy score between 7 and 27, resulting in groups with comparable overall levels of chromosomal instability. After filtering, the distribution of other chromosome-arm alterations between loss and neutral groups was markedly reduced, minimizing confounding in downstream comparisons.

To identify chromosome-arm-specific vulnerability genes, cancer cell lines were grouped into chromosome-arm-loss and neutral categories. Differences in genetic dependencies between groups were assessed using the Wilcoxon-Mann-Whitney test. A gene was classified as a significant chromosome-loss-associated dependency if all of the following criteria were met: (i) the difference in median dependency scores (chromosome neutral - chromosome loss) exceeded 0.1; (ii) the median dependency score in the chromosome-loss group was below - 0.3; and (iii) the *P* value was less than 0.01. Paralog-dependency annotations were assigned using paralog information obtained from Ensembl^106^. Interaction-dependency annotations were derived from the published protein-protein interaction database STRING^107^.

### Paralog pair annotation

Paralog information was obtained from Ensembl^106^. For comparative analyses, paralog pairs were included only if they exhibited an identity score of at least 25% and belonged to paralog families with fewer than 20 members.

### Gene Ontology (GO) enrichment analysis

GO enrichment analysis was performed using the ClusterProfiler R package (v4.14.6)^108^. Enriched biological processes, molecular functions, and cellular components were identified using default parameters unless otherwise specified. All downstream data processing, visualization, and plotting were conducted in R using custom scripts built with the tidyverse package (v2.0.0)^101^.

### Gene evolutionary conservation analysis

Gene evolutionary conservation was assessed using PhyloP scores across the exonic regions of each paralog gene^109^. Exonic PhyloP scores were compared between paralogs in significant dependency pairs and those in non-significant pairs.

### Subcellular co-localization analysis

Subcellular localization annotations were obtained from the Human Protein Atlas^110^. For each paralog pair, we determined whether the two proteins shared at least one annotated subcellular location. The frequency of shared localization was then compared between significant dependency pairs and non-significant paralog pairs.

### Paralog vulnerability validation in shRNA screens

To assess whether the paralog dependencies identified in the CRISPR analysis were also observed in RNAi screens, we analyzed DepMap DEMETER2 dependency scores for the ten most significant paralog pairs^111^. We repeated the correlation analysis using the same cutoffs applied in the CRISPR analysis.

### Paralog dosage compensation

#### Analysis of chromatin accessibility at paralog genes

Preprocessed ATAC-seq bigWig signal files were obtained from the published TCGA pan-cancer dataset, which comprised 410 tumor samples spanning 23 cancer types^36^. Chromosome loss events in TCGA were obtained from ref^9^. ATAC-seq signal surrounding the transcription start sites (TSS) of significant and non-significant paralog genes was extracted using deepTools “computeMatrix reference-point”, with 3-kb upstream and downstream windows and a bin size of 50 bp^112^. For each chromosome-loss event, chromatin accessibility at significant and non-significant paralog genes was compared between tumors with chromosome loss and those with neutral chromosome status. Analyses were restricted to tumors classified as diploid to limit the effects of whole-genome doubling. To account for gene dosage, ATAC-seq signal was divided by the chromosome copy number: one for chromosome loss and two for neutral chromosome status. To test whether chromosome loss altered chromatin accessibility differently at significant and non-significant paralog genes, we quantified ATAC-seq signal within ∼300 bp upstream and downstream of each transcription start site. For each group, the accessibility change was calculated by comparing chromosome-loss and chromosome-neutral tumors. We then used a one-sided permutation test with 10,000 permutations to assess whether the accessibility change was greater for non-significant than for significant paralog genes.

#### Analysis of H3K27ac histone modification at paralog genes

Preprocessed H3K27ac HiChIP bigWig signal files were obtained from a published pan-cancer dataset comprising 69 tumor samples across 15 cancer types^37^. H3K27ac signal surrounding the transcription start sites of significant and non-significant paralog genes was extracted using deepTools “computeMatrix reference-point”, with 3-kb upstream and downstream windows and a bin size of 50 bp^112^. For each chromosome-loss event, H3K27ac signal was compared between tumors with chromosome loss and those with neutral chromosome status, separately for significant and non-significant paralog genes. Analyses were restricted to tumors classified as diploid and to chromosome-loss events represented by at least five samples. To account for gene dosage, H3K27ac signal was divided by the chromosome copy number: one for chromosome loss and two for neutral chromosome status. To test whether chromosome loss was associated with different changes in H3K27ac signal at significant and non-significant paralog genes, we quantified signal within ∼300 bp upstream and downstream of each TSS. For each group, the change in signal was calculated by comparing chromosome-loss and chromosome-neutral tumors. A one-sided permutation test with 10,000 permutations was used to assess whether the change was greater for non-significant than for significant paralog genes.

#### Analysis of dosage compensation in significant and non-significant paralogs

To assess dosage compensation in paralogs classified as significant or non-significant in the dependency analysis, we analyzed CCLE and TCGA gene expression data represented as log₂(TPM + pseudocount). For each paralog, median expression was calculated separately across samples with loss of the corresponding chromosome arm and samples with a copy-number-neutral chromosome arm. Relative expression was defined as the median log-transformed expression in the chromosome-arm-loss group minus that in the copy-number-neutral group. Relative expression distributions were compared between significant and non-significant paralogs using a two-sided Wilcoxon rank-sum test.

#### Analysis of dosage compensation in significant and non-significant paralogs in isogenic cell lines

To assess dosage compensation following chromosome-arm loss, we analyzed gene expression datasets from isogenic cell lines: GSE68042 (8p loss)^113^, GSE150686 (19p loss)^114^, GSE220320 (8p loss)^39^, GSE293644 (3p and 8p loss)^115^, and Shih et al. (8p loss)^29^. HLF samples from GSE220320 were excluded because we did not detect an overall reduction in the expression of genes on the nominally lost chromosome arm. Within each dataset, genes were retained if their expression value was greater than 0 in at least one sample, excluding genes with no detectable expression across all samples. For significant and non-significant paralog genes located on the corresponding lost chromosome arm, expression in chromosome-arm-loss cells was normalized to that in matched chromosome-neutral isogenic controls. Relative expression was compared between significant and non-significant paralogs using a two-sided Wilcoxon rank-sum test.

### Analysis of dependencies associated with loss of the Y chromosome

#### Feature matrix

Copy-number and mutation features were restricted to the MSK-IMPACT panel (*n* = 505 genes)^116^. For copy number, genes in the pseudo-autosomal regions or on chromosome Y were excluded to avoid circularity with LOY status, leaving 500 genes encoded as gain, loss or wild type relative to sample ploidy. For mutations, calls were restricted to protein-coding variants with a predicted functional consequence, and genes mutated in fewer than 5% of cell lines were removed, leaving 62 genes encoded as mutant or wild type. Lineages with fewer than ten cell lines in either the Y-WT or LOY group were collapsed into a single “Other” category. MSI status was defined from broad MSI scores, with samples classified as MSI-high at a score ≥ 7. Whole-genome doubling status and aneuploidy score were excluded, as these annotations were unavailable for a substantial proportion of the cohort. After complete-case filtering, the final dataset comprised *n* = 622 cell lines and *p* = 566 features.

#### Linear regression

Univariate associations were assessed using linear models with DDX3X gene effect as the response. A base model containing LOY status alone established the overall association between chromosome Y loss and DDX3X dependency. Additive models (dependency ∼ LOY + feature) tested whether each feature improved model fit beyond LOY alone, and interaction models (dependency ∼ LOY × feature) tested whether the association between a feature and DDX3X dependency differed by LOY status.

#### Elastic net regression

Multivariable modelling followed the approach of Garnett et al. (2012), using glmnet with DDX3X gene effect (Chronos) as the response^117^. The mixing parameter α was fixed at 0.5, giving equal weight to the L1 and L2 penalties. This differs from Garnett et al., who optimized both penalty parameters, and was informed by a preliminary sweep across α from 0.1 to 1.0 in increments of 0.1: cross-validated mean squared error decreased from α = 0.1 to α ≈ 0.3 and was essentially flat thereafter, varying by less than 0.3% across the range 0.3-1.0. Fixing α therefore incurred negligible cost in model fit while holding sparsity constant across iterations. Within each iteration the penalty λ was selected by ten-fold cross-validation minimizing mean squared error, and a final model was refitted on the full dataset at the selected λ.

This procedure was repeated over 100 iterations, each with an independently drawn random fold assignment, to assess the stability of feature selection under repeated cross-validation. For each feature we recorded the selection frequency (*f*), the proportion of iterations in which the coefficient was non-zero, and the effect size, defined as the coefficient multiplied by the standard deviation of that feature across the panel and averaged over all 100 iterations, including those in which the feature was not selected. Effect-size standardization accounts for the differing scales on which genomic features are measured. Features were considered highly significant when both the selection frequency (*f* > 0.48) and the mean effect size (outside -0.0099 to 0.0095) exceeded the corresponding panel mean by more than two standard deviations, yielding 19 features. Thresholds were derived empirically from the distributions observed in this dataset rather than adopted from published values.

### Comparing DDX3X with published aneuploidy and clinical vulnerabilities

To compare the strength of the DDX3X dependency associated with loss of the Y chromosome with previously reported aneuploidy-associated therapeutic vulnerabilities, we reanalyzed the published targets using DepMap dependency data. A high-confidence LOY group was defined by the absence of detectable copy number across 25 Y-linked genes and the absence of detectable DDX3Y expression. A high-confidence Y-positive group was defined by detectable Y-linked gene copy number and DDX3Y expression. For analyses of individual chromosome-arm alterations, cell lines were filtered by aneuploidy score using the criteria applied in the chromosome-loss dependency analysis. To assess dependencies associated with overall aneuploidy, cell lines in the top and bottom quartiles of aneuploidy score were classified as high and low aneuploidy, respectively. Dependency scores were compared between biomarker-positive and biomarker-negative groups using a Wilcoxon rank-sum test.

To compare the strength of the DDX3X dependency associated with loss of the Y chromosome with previously reported dependencies linked to clinical biomarkers, we reanalyzed the published targets using DepMap dependency data. Mutation and microsatellite instability (MSI) status were classified as described in the “Feature matrix” section. Dependency scores were compared between biomarker-positive and biomarker-negative cell lines using a Wilcoxon rank-sum test.

### Statistical Analysis

All experiments were performed a minimum of two times with a minimum of three independent measurements. All statistical analysis was performed with R 4.4.2, or Python 3.10. All data are represented as mean ± standard error.

## Code availability

Code generated to perform this analysis is available at https://github.com/sheltzer-lab/Paralog_Dependency.

## Data availability

The raw proteomics data have been deposited to the ProteomeXchange Consortium via the MassIVE partner repository, with the dataset identifier MSV000103695.

## Acknowledgments

Research in the Sheltzer Lab is supported by NIH grant R01CA276666, a Melanoma Research Alliance grant (#1576215), a Hevolution/AFAR grant, an Andrew McDonough B+ Foundation grant, a Blavatnik Fund award, a grant from the Colorectal Cancer Alliance, and an American Cancer Society Mission Boost grant. I.F. was supported by the Propel Postdoctoral Scholars Program, Stanford School of Medicine. We thank the Stanford Veterinary Service Center for assistance with the mouse experiments. This work utilized Stanford Cancer Institute Shared Resources, which are supported in part by NIH P30 CA124435. S.R. acknowledges funding from Deutsche Forschungsgemeinschaft (DFG, German Research Foundation): project-ID 469332207 and project-ID 493697503. For this research, A.M.T. acknowledges NIH Grant R35GM147287.

## Supplemental figure legends

## Supplemental figure legends

**Supplementary Figure 1.**
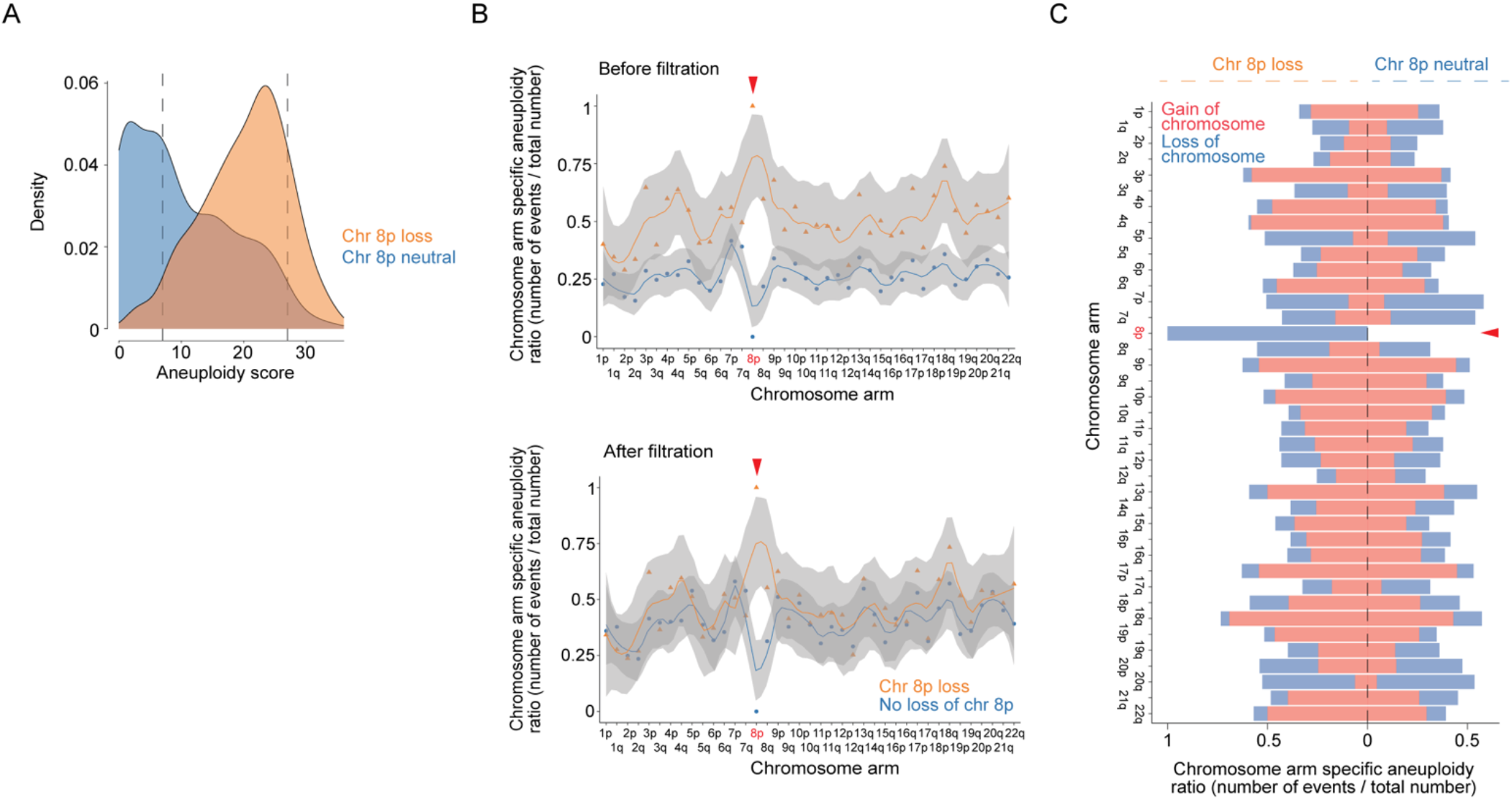
Aneuploidy score correction to identify dependencies associated with specific chromosome loss events. (A) Distributions of aneuploidy scores in cancer cell lines with chromosome 8p loss (orange) or neutral 8p copy-number status (blue). (B) Frequencies of chromosome-arm aneuploidy events in the two groups before (top) and after (bottom) aneuploidy-score filtering. Each frequency represents the number of cell lines with an event affecting the indicated arm divided by the total number of cell lines in the group. Arrowheads mark chromosome 8p. (C) Frequencies of chromosome-arm gains (red) and losses (blue) after aneuploidy-score filtering, shown for cell lines with chromosome 8p loss (left) or neutral 8p copy-number status (right). The arrowhead marks chromosome 8p.

**Supplementary Figure 2.**
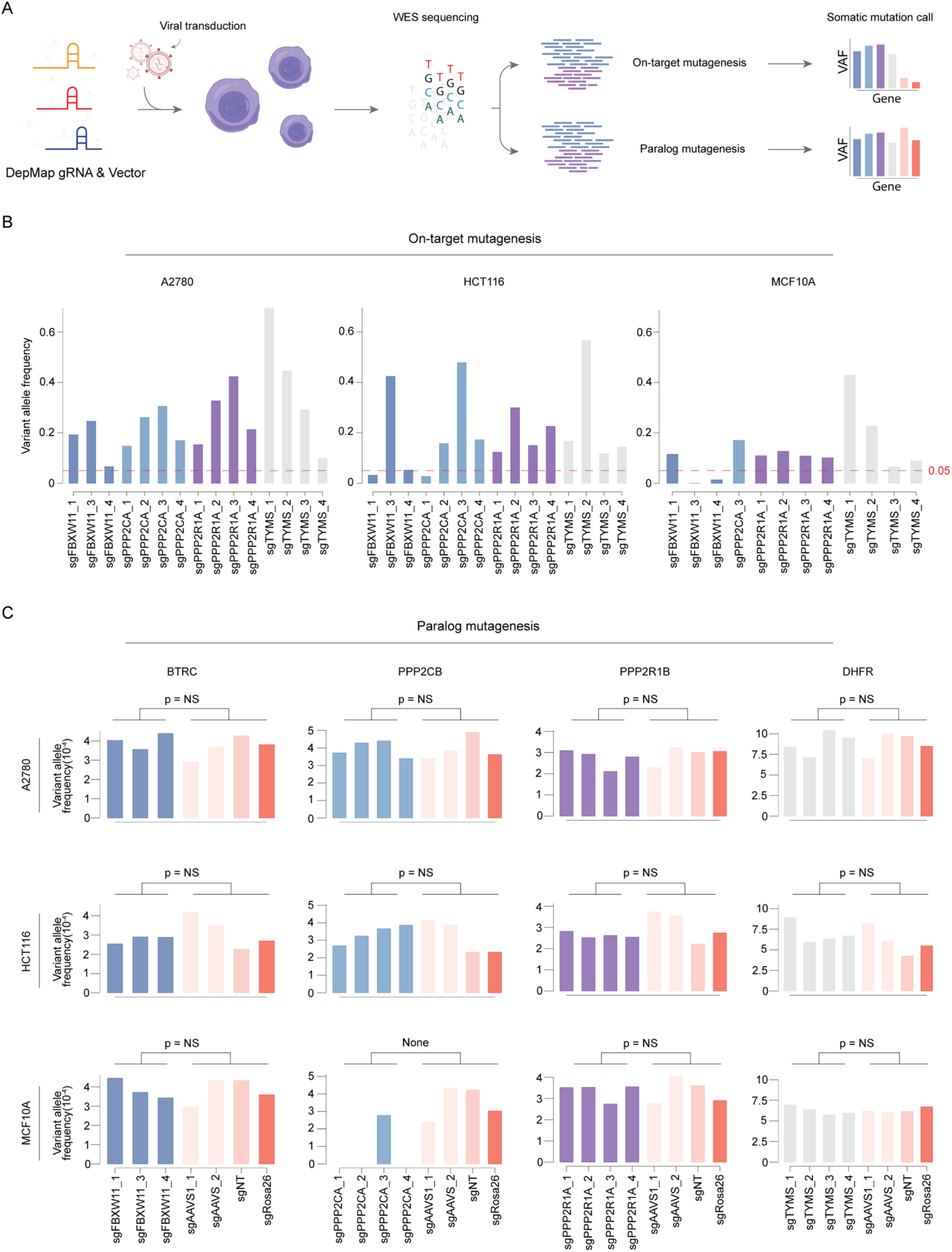
No evidence for off-target CRISPR-induced mutations in paralogs of a targeted gene. (A) Workflow for assessing on-target and off-target paralog mutagenesis by whole-exome sequencing after transduction with the indicated DepMap gRNAs. (B) Variant allele frequencies at *FBXW11*, *PPP2CA*, *PPP2R1A* and *TYMS* loci in A2780, HCT116, and MCF10A cells following gRNA transduction and selection. Mutations were assessed within 10 bp upstream and downstream of the CRISPR cut site. Dashed lines indicate a variant allele frequency of 0.05. (C) Variant allele frequencies in *BTRC*, *PPP2CB*, *PPP2R1B* and *DHFR* after targeting their respective paralogs, *FBXW11*, *PPP2CA*, *PPP2R1A* and *TYMS*, in A2780, HCT116, and MCF10A cells. Cells were transduced with gRNAs targeting *AAVS1*, *Rosa26*, and non-targeting gRNAs as controls.

**Supplementary Figure 3.**
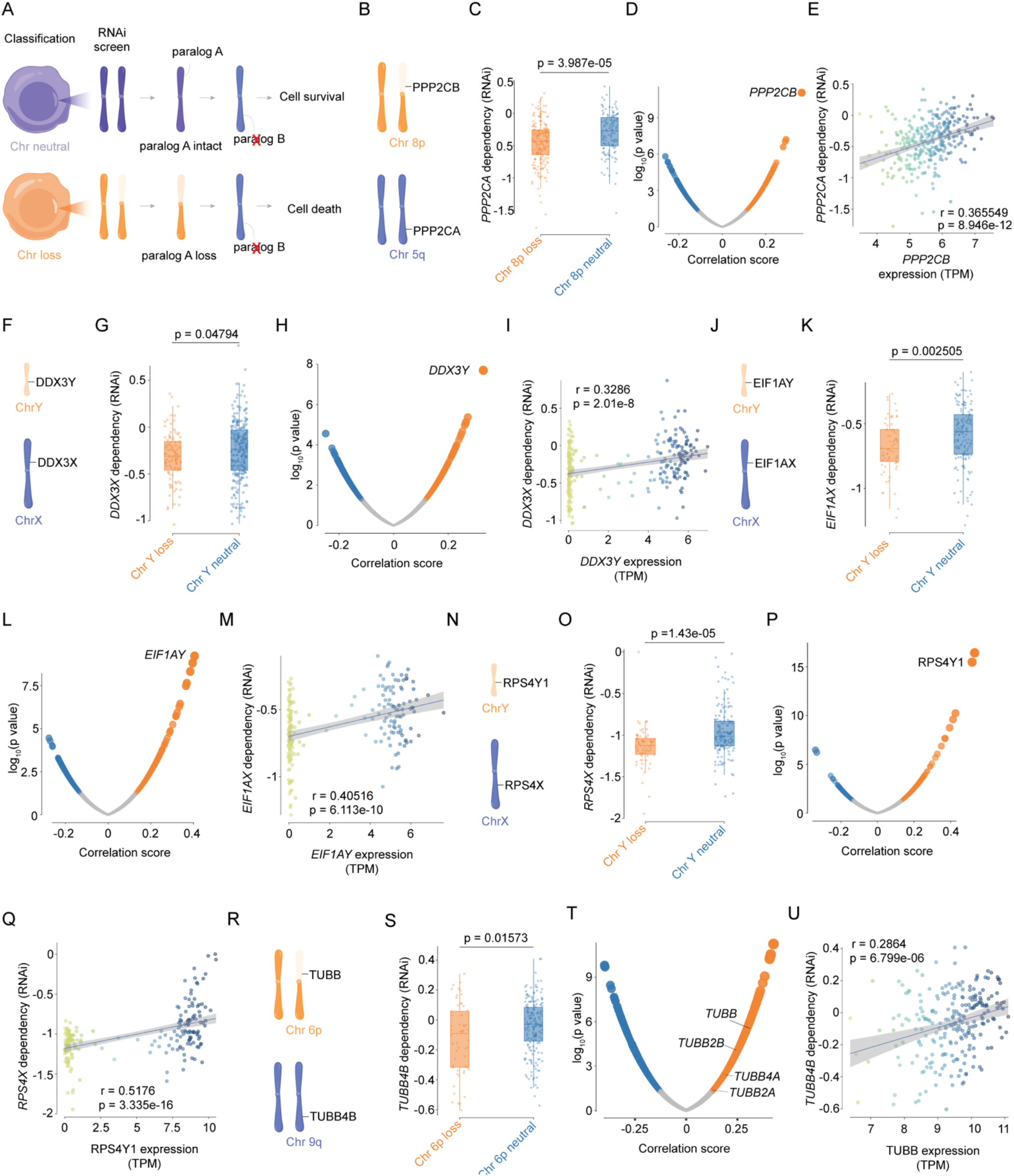
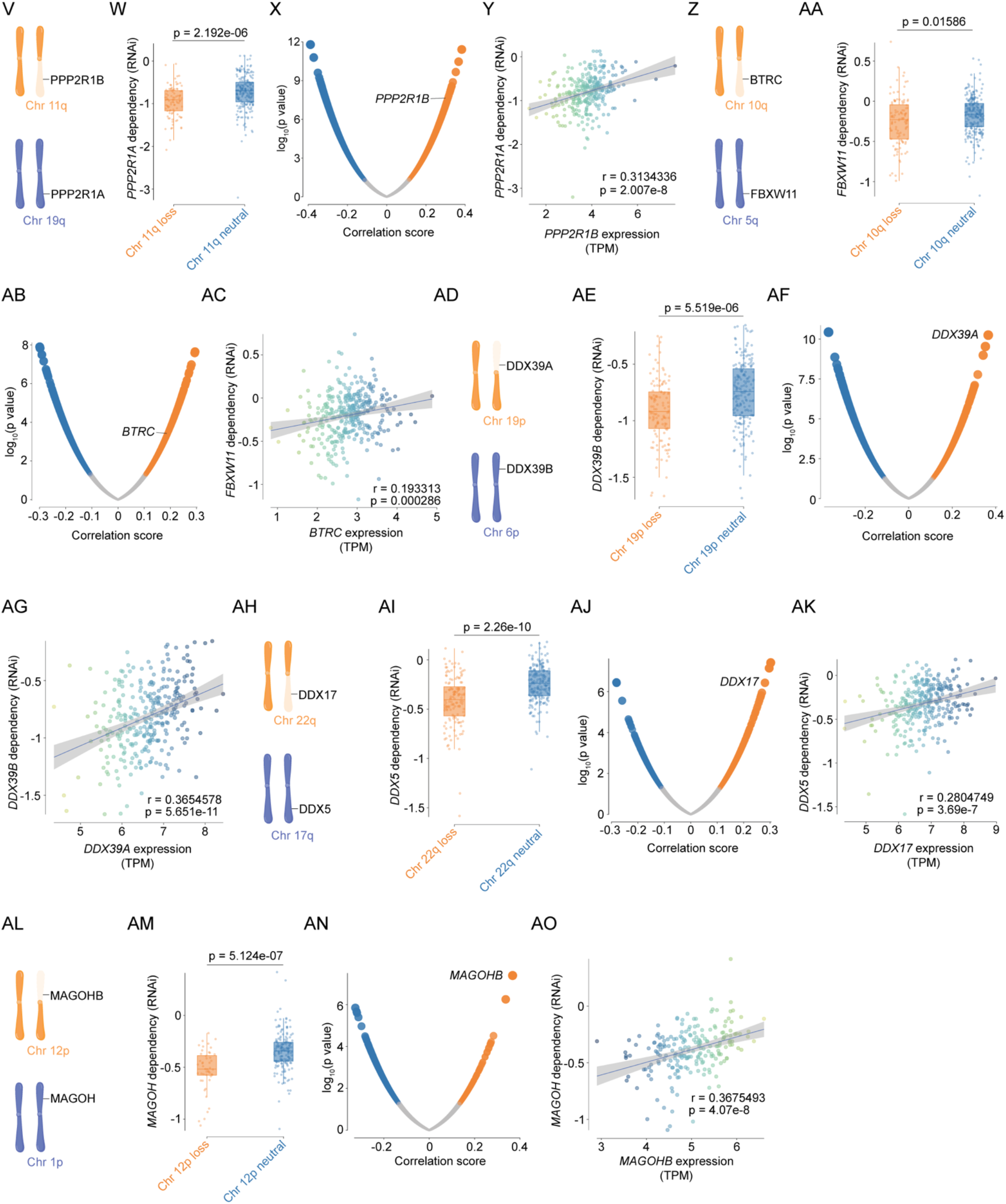
Validation of chromosome loss-associated paralog vulnerabilities in RNAi screens. (A) Workflow for evaluating chromosome-loss-associated paralog dependencies in DepMap RNAi screens. (B) Chromosomal locations of *PPP2CB* on 8p and *PPP2CA* on 5q. (C) *PPP2CA* RNAi dependency scores in cell lines with chromosome 8p loss or neutral 8p copy-number status. (D) Genome-wide correlations between gene expression and *PPP2CA* RNAi dependency scores, with *PPP2CB* labeled. (E) Relationship between *PPP2CB* expression and *PPP2CA* RNAi dependency scores. (F) Chromosomal locations of *DDX3Y* on the Y chromosome and *DDX3X* on the X chromosome. (G) *DDX3X* RNAi dependency scores in cell lines with Y chromosome loss or an intact Y chromosome. (H) Genome-wide correlations between gene expression and *DDX3X* RNAi dependency scores, with *DDX3Y* labeled. (I) Relationship between *DDX3Y* expression and *DDX3X* RNAi dependency scores. (J) Chromosomal locations of *EIF1AY* on the Y chromosome and *EIF1AX* on the X chromosome. (K) *EIF1AX* RNAi dependency scores in cell lines with Y chromosome loss or an intact Y chromosome. (L) Genome-wide correlations between gene expression and *EIF1AX* RNAi dependency scores, with *EIF1AY* labeled. (M) Relationship between *EIF1AY* expression and *EIF1AX* RNAi dependency scores. (N) Chromosomal locations of *RPS4Y1* on the Y chromosome and *RPS4X* on the X chromosome. (O) *RPS4X* RNAi dependency scores in cell lines with Y chromosome loss or an intact Y chromosome. (P) Genome-wide correlations between gene expression and *RPS4X* RNAi dependency scores, with *RPS4Y1* labeled. (Q) Relationship between *RPS4Y1* expression and *RPS4X* RNAi dependency scores. (R) Chromosomal locations of *TUBB* on 6p and *TUBB4B* on 9q. (S) *TUBB4B* RNAi dependency scores in cell lines with chromosome 6p loss or neutral 6p copy-number status. (T) Genome-wide correlations between gene expression and *TUBB4B* RNAi dependency scores, with *TUBB*, *TUBB2B*, *TUBB4A* and *TUBB2A* labeled. (U) Relationship between *TUBB* expression and *TUBB4B* RNAi dependency scores. (V) Chromosomal locations of *PPP2R1B* on 11q and *PPP2R1A* on 19q. (W) *PPP2R1A* RNAi dependency scores in cell lines with chromosome 11q loss or neutral 11q copy-number status. (X) Genome-wide correlations between gene expression and *PPP2R1A* RNAi dependency scores, with *PPP2R1B* labeled. (Y) Relationship between *PPP2R1B* expression and *PPP2R1A* RNAi dependency scores. (Z) Chromosomal locations of *BTRC* on 10q and *FBXW11* on 5q. (AA) *FBXW11* RNAi dependency scores in cell lines with chromosome 10q loss or neutral 10q copy-number status. (AB) Genome-wide correlations between gene expression and *FBXW11* RNAi dependency scores, with *BTRC* labeled. (AC) Relationship between *BTRC* expression and *FBXW11* RNAi dependency scores. (AD) Chromosomal locations of *DDX39A* on 19p and *DDX39B* on 6p. (AE) *DDX39B* RNAi dependency scores in cell lines with chromosome 19p loss or neutral 19p copy-number status. (AF) Genome-wide correlations between gene expression and *DDX39B* RNAi dependency scores, with *DDX39A*labeled. (AG) Relationship between *DDX39A* expression and *DDX39B* RNAi dependency scores. (AH) Chromosomal locations of *DDX17* on 22q and *DDX5* on 17q. (AI) *DDX5* RNAi dependency scores in cell lines with chromosome 22q loss or neutral 22q copy-number status. (AJ) Genome-wide correlations between gene expression and *DDX5* RNAi dependency scores, with *DDX17* labeled. (AK) Relationship between *DDX17* expression and *DDX5* RNAi dependency scores. (AL) Chromosomal locations of *MAGOHB* on 12p and *MAGOH* on 1p. (AM) *MAGOH* RNAi dependency scores in cell lines with chromosome 12p loss or neutral 12p copy-number status. (AN) Genome-wide correlations between gene expression and *MAGOH* RNAi dependency scores, with *MAGOHB* labeled. (AO) Relationship between *MAGOHB* expression and *MAGOH* RNAi dependency scores. In the dependency comparisons, chromosome-loss and chromosome-neutral groups are shown in orange and blue, respectively. Expression–dependency scatter plots show individual cell lines, fitted regression lines, correlation coefficients and P values.

**Supplementary Figure 4.**
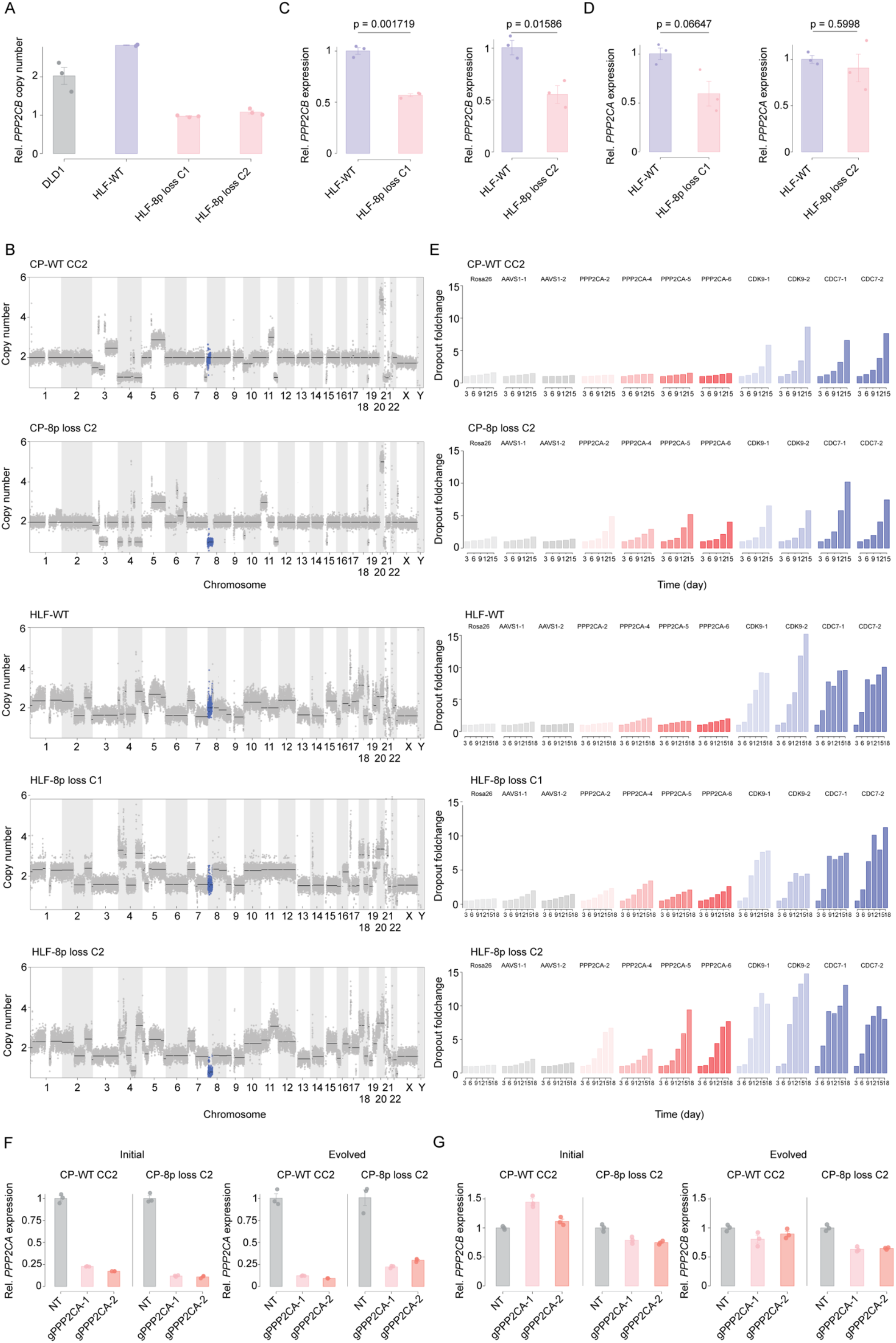
Characterization of isogenic 8p-loss models and their responses to *PPP2CA* suppression. (A) Relative *PPP2CB* copy number in DLD1 cells, HLF-WT cells and two HLF 8p-loss clones. DLD1 cells served as a diploid control. (B) Genome-wide copy-number profiles of CP-WT CC2, CP-8p loss C2, HLF-WT, HLF-8p loss C1 and HLF-8p loss C2 cells. Chromosome 8p is highlighted in blue. (C) Relative *PPP2CB* expression in HLF-WT cells and each of the two HLF 8p-loss clones. (D) Relative *PPP2CA* expression in HLF-WT cells and each of the two HLF 8p-loss clones. (E) CRISPR competition assays in the cell lines shown in B, measuring dropout at the indicated time points after targeting with four independent *PPP2CA* gRNAs. *Rosa26* and *AAVS1* guides served as negative controls, and *CDK9* and *CDC7* guides served as pan-essential positive controls. (F) Relative *PPP2CA* expression in CP-WT CC2 and CP-8p loss C2 cells carrying a non-targeting guide (NT) or two *PPP2CA*-targeting CRISPRi guides, measured at the initial and evolved time points. (G) Relative *PPP2CB* expression in the same cells from (F) at the initial and evolved time points.

**Supplementary Figure 5.**
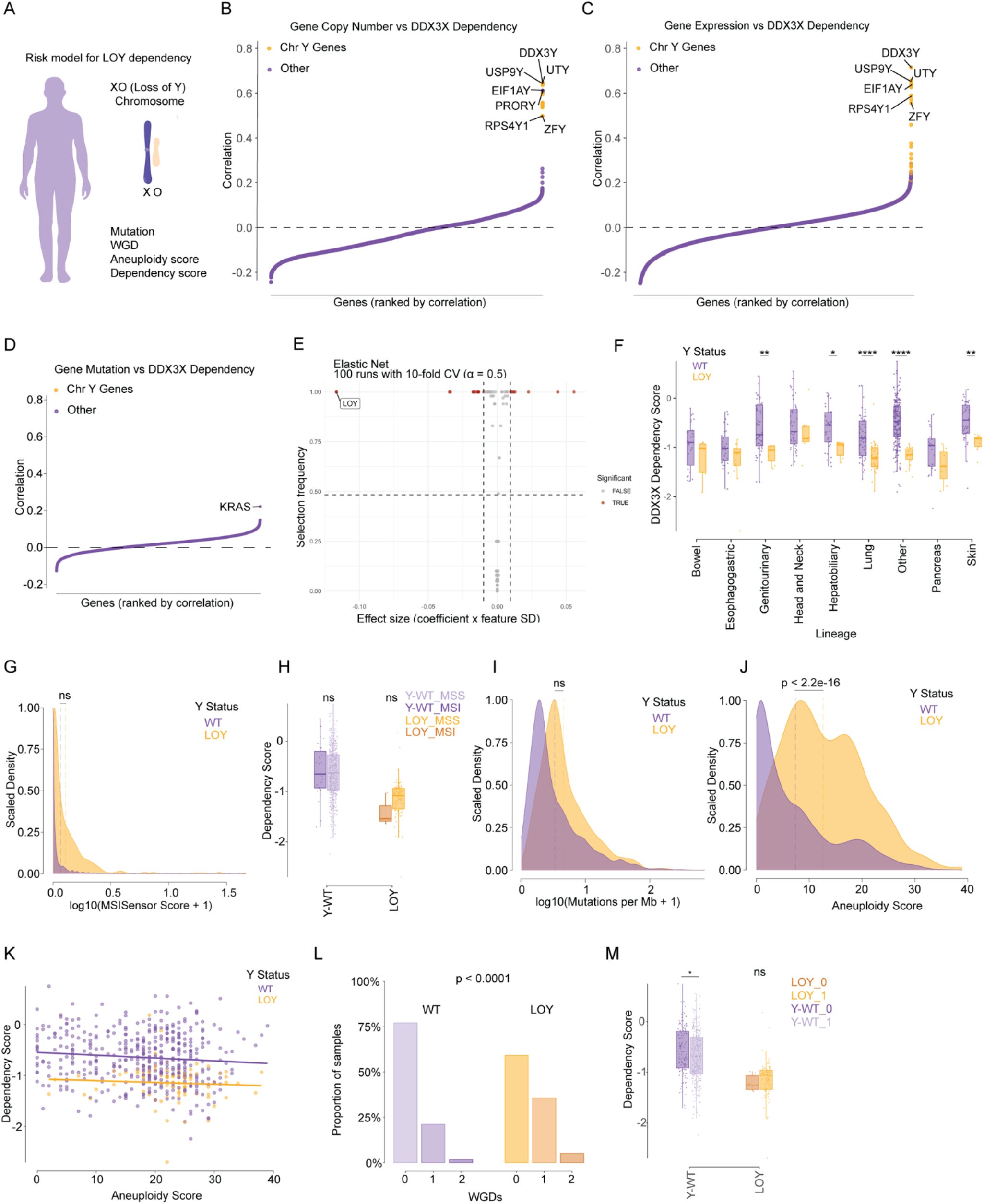
Genomic correlates of DDX3X dependency across cancer cell lines. (A) Overview of the analysis of Y chromosome status and *DDX3X* dependency in male cancer cell lines. (B) Pearson correlations between gene copy number and *DDX3X* dependency scores. Genes encoded on the Y chromosome are highlighted in orange. (C) Pearson correlations between gene expression and *DDX3X* dependency scores. Genes encoded on the Y chromosome are highlighted in orange. (D) Pearson correlations between gene mutations and *DDX3X* dependency scores. (E) Effect sizes and selection frequencies of predictors in an elastic net model of *DDX3X* dependency, including Y chromosome status, lineage and genomic alterations. Results summarize 100 runs with 10-fold cross-validation and α = 0.5. (F) *DDX3X* dependency scores in cancer cell lines with an intact Y chromosome (WT) or Y chromosome loss (LOY), stratified by lineage. (G) Distributions of microsatellite instability scores in cancer cell lines with an intact Y chromosome or LOY. (H) *DDX3X* dependency scores in microsatellite-stable (MSS) and microsatellite-unstable (MSI) cancer cell lines, stratified by Y chromosome status. (I) Distributions of mutation burden in cancer cell lines with an intact Y chromosome or LOY. (J) Distributions of aneuploidy scores in cancer cell lines with an intact Y chromosome or LOY. (K) Relationship between aneuploidy score and *DDX3X* dependency in cancer cell lines with an intact Y chromosome or LOY. Regression lines are shown separately for each group. (L) Proportions of cancer cell lines with zero, one or two whole-genome doubling (WGD) events, stratified by Y chromosome status. (M) *DDX3X* dependency scores in cancer cell lines with or without WGD, stratified by Y chromosome status.

**Supplementary Figure 6.**
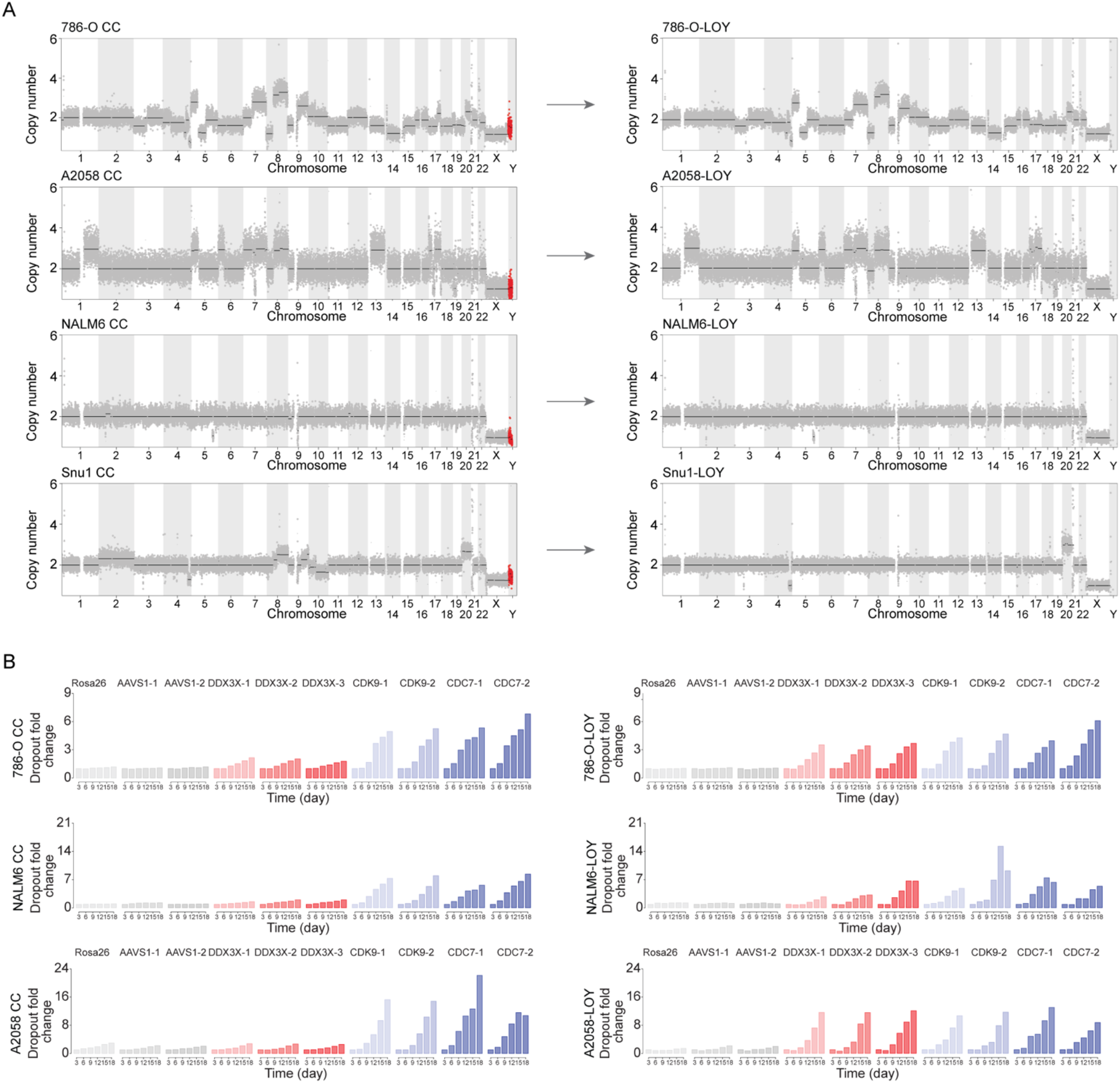
Loss of Y causes a specific dependency on *DDX3X* in additional isogenic models. (A) Genome-wide copy-number profiles of isogenic wild-type and LOY derivatives of 786-O, A2058, NALM6 and Snu1 cells. The Y chromosome is highlighted in red. (B) CRISPR competition assays in isogenic wild-type and LOY derivatives of 786-O, NALM6 and A2058 cells, showing dropout over 18 days following targeting with three independent *DDX3X* gRNAs. *Rosa26* and *AAVS1* guides served as negative controls, and *CDK9* and *CDC7* guides served as pan-essential positive controls.

**Supplementary Figure 7.**
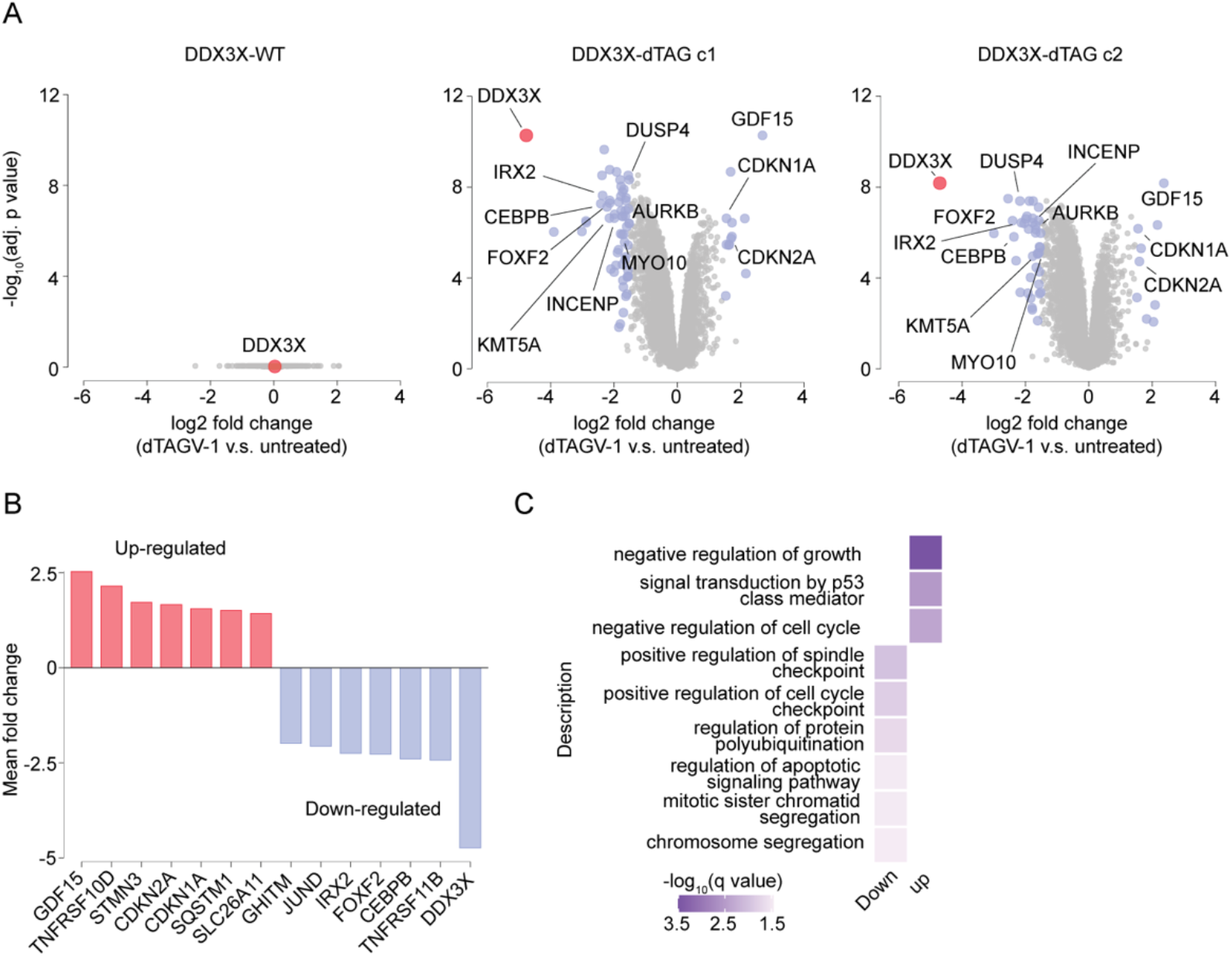
Quantitative analysis of the proteome following DDX3X ablation. (A) Volcano plots showing changes in protein abundance following treatment with 50 nM dTAGV-1 for 24 hours in SNU1-LOY cells expressing untagged DDX3X (DDX3X-WT) or two independent DDX3X-dTAG clones (c1 and c2). DDX3X is highlighted in red, and the complete results are included in Table S6A. (B) Log₂ fold changes in the abundance of selected proteins following DDX3X degradation. Proteins with increased abundance are shown in red, and proteins with decreased abundance are shown in purple. (C) Gene Ontology enrichment analysis of proteins with increased or decreased abundance following DDX3X degradation. Selected biological processes are shown, with color intensity indicating enrichment significance. Complete results are presented in Table S6B-C.

